# Phytoliths to apPROXYmate Maya landscapes over early Anthropocene : case of Naachtun tropical wetland

**DOI:** 10.1101/2025.03.05.641636

**Authors:** M. Testé, C. Castanet, A. Garnier, N. Limondin-Lozouet, L. Purdue, E. Lemonnier, S. Kerdreux, P. Non-dédéo

## Abstract

Reconstitutions of past anthromes in the tropical karstic environments of the Central Maya lowlands (CMLs) are generally restrained by bioindicators such as pollen in the rare lakes and perennial wetland systems and charcoals. This original work mobilizes phytoliths and, to a lesser extent, other bioindicators such as molluscs and diatoms. It contributes to demonstrate of these fossils’ potential to participate in the reconstruction of the Maya society-environment interactions, in the CMLs Elevated Interior Region. We studied swamp and lake sediment archives of a karst polje (nammed *bajo El Infierno*), located north of the regional Maya city of Naachtun. These bioindicators were studied on three sediment series covering the last 3500 years, during which this region was settled by the Maya. This multiproxy and multi-local approache allowed to propose an environnemental history for this wetland. It shows multi-millennial anthropogenic impacts in the *bajo*. The local opening of the swamp forest suggests the occupation of this *bajo* during the early Preclassic. The bioindicators also provide information on the variability of lake conditions, marked in particular by perennial shallow lake dynamics during the middle Preclassic, which provided a resource for the Maya. The direct trace of maize agriculture and pervasive herbaceous formations through the *bajo* are finally noted during the late Preclassic. Demographical and political disruptions occurred to Naachtun during the Preclassic-Classic transition, simultaneous to regional droughts. The Classic is marked by the presence of an environmental mosaic which may indicate a change in *bajo* land use. This landscape diversity demonstrates a new time the need to carry out multi-local paleoecological approaches to understand the society-environment interactions. Phytoliths, by their siliceous mineralization and local expression, seem appropriate to become an essential tool for these approaches in the CMLs.

## 1. Introduction

The use of phytoliths to reconstruct tropical palaeoecosystems has been successful in recent years, particularly in Latin America (Piperno et al., 2007; Iriarte et al., 2010; Watling et al., 2017). However, palaeoecological studies in the Maya Lowlands are mainly limited to the use of pollen, which is rare in these tropical karst regions (Islebe et al., 1996; Leyden, 2002; Wahl et al., 2006; Carozza et al., 2007; Carrillo-Bastos et al., 2012). Researches with phytoliths are poorly developed and rarely concerned palaeoenvironments. They mainly focus on the archaeological signature of human domestic practices (Cummings & Magenis, 1997; Bozarth & Guderjan, 2004; Abramiuk et al., 2011). Recent studies on wetlands at Maya archaeological sites have used phytoliths, but these only offer limited assemblages (Beach et al., 2009; Krause et al., 2019, 2021).

This reluctance to use phytoliths in palaeoenvironmental studies may explain by, supposedly, their intrinsic taxonomic biases that limit reflection based on indicator species (Rovner, 1971). Phytolithists then characterize environments with assemblages of morphotypes (Neumann et al., 2009) and build modern references to study plant communities’ signature (Dickau et al., 2013; Watling et al., 2016). Such a reference frame has recently been developed on the modern ecosystems of the archaeological site of Naachtun, located in the extreme north of Guatemala (Testé et al., 2020). This work was initiated because of the sediments’ high phytolith content in the seasonal drained swamp area (locally named *bajo*). Besides, the deposits of this *bajo* recorded many other bioindicators that can be studied in addition to the phytoliths: sponge spicules, diatoms, and fossil shells of gastropods.

Naachtun is a site in the Petén with definite archaeological interest. Its political and architectural center developed in the late Preclassic (150 CE) when most large neighboring cities, such as Nakbe and El Mirador, were abandoned (Wahl et al., 2007, Hiquet, 2020). Naachtun has been a regional capital for a millennium and played an essential role in the central Lowlands’ political history through its alliances with Tikal and Calakmul’s regional powers. The latter and many cities in the Central Maya Lowlands (CMLs) are abandoned at the terminal Classic, even as Naachtun’s population remains stable. The city is, in turn, abandoned around 950 CE (Sion, 2016). This history of Naachtun, disharmonious with regional dynamics, makes it an appropriate study site to understand the Maya populations’ adapt-ability to their territory’s particularities. This requires an understanding of agricultural practices and the management of resources such as water and soil. Recent LiDAR remote sensing work has highlighted a dense network of anthropogenic structures (canals, reservoirs, terraces) in the city’s surroundings, particularly in the wetlands and the *bajo* (Canuto et al., 2018). They reinforce the interest in these peripheral areas of the Maya cities and their possible agricultural function.

If this work’s primary intention is to reconstitute past vegetation in an issue of society-environment interactions, there are underlying questions that drive its development. A first concern is the capacity of phytoliths to characterize past environments, with interest in their accuracy. While the modern reference of Naachtun has made it possible to distinguish modern environments, uncertainties about ancient plant communities are still significant. It remains to be determined whether the fossil assemblages will make it possible to find these modern environments and interpret environments that are not referenced in Naachtun today, such as anthropized environments. This questioning also concerns the possibility of identifying phytoliths of cultivars in agrarian contexts far from the site since so far they have only been observed regionally in archaeological contexts (Cummings & Magenis, 1997; Bozarth & Guderjan, 2004). The parallel study of several bioindicators aims at coupling ecological signals to allow a better palaeoenvironmental reconstruction. However, concentrations of diatoms and sponge spicules have only been assessed in *bajo* ecosystems in Naachtun (Testé et al., 2020). The suspected presence of former lake conditions not referenced in the current one could make the results on these bioindicators uninterpretable. Finally, despite a well-established taxonomy, the autecology of freshwater gastropods of the Petén is poorly known. Only rare ecological reference frames present the ecological valences of freshwater species of the lowlands (Goodrich & Van der Schalie, 1937). The study of fossil mollusk assemblages in the *bajo* sediments is therefore aimed at verifying whether a possible hydrological or botanical signal can confirm that presented by the phytoliths and siliceous bioindicators.

This study rest upon one auger drilling and two boreholes pit in the clayey sediments of the *bajo* zone, north of the site. Each of these series was studied by way of 20 samples, in which were evaluated phytoliths assemblages, concentrations of sponge spicules, and diatoms. In parallel with the phytoliths study, the malacological groups of two deep boreholes, 40 samples in total, were studied and compared with the microsilica bioindicator data. This paper proceeds with a synoptic presentation of the stratigraphical, sedimentary and chronological contexts as described in Castanet et al. (2022). The results on assemblages of siliceous phytoliths and bioindicators are presented and interpreted according to different approaches. Malacological groups are shown in the same way. Finally, this study proposes a comparative reading of the naturalist data obtained with the site’s archaeological data. This unprecedented data correlation aims better to understand the society-environment interactions in these particular tropical ecosystems.

## 2. Study area

### 2.1. The archaeological site of Naachtun

The site of Naachtun (Fig.1), in the extreme north of Guatemala, has been known to the chicleros (former sap collectors, *Manilkara zapota*) since at least 1916, when it was reported by one of them, Alfonso Ovando (Morley, 1938). The site was partially studied during the first part of the 20th century (Lundell, 1932; Morley, 1938) before renewed interest in these ruins from the 2000s (Reese-Taylor et al., 2005). Since 2010, the Franco-Guatemalan program (CNRS/Université Paris I Panthéon Sorbonne - San Carlos University of Guatemala) has brought together teams of archaeologists and palaeoenvironmentalists. It has enabled 200 Ha of the site to be studied (Nondédéo et al., 2014), i.e., 150 Ha more than in previous projects, and has recently been LIDAR coverage (Canuto et al., 2018).

**Figure 1-.**
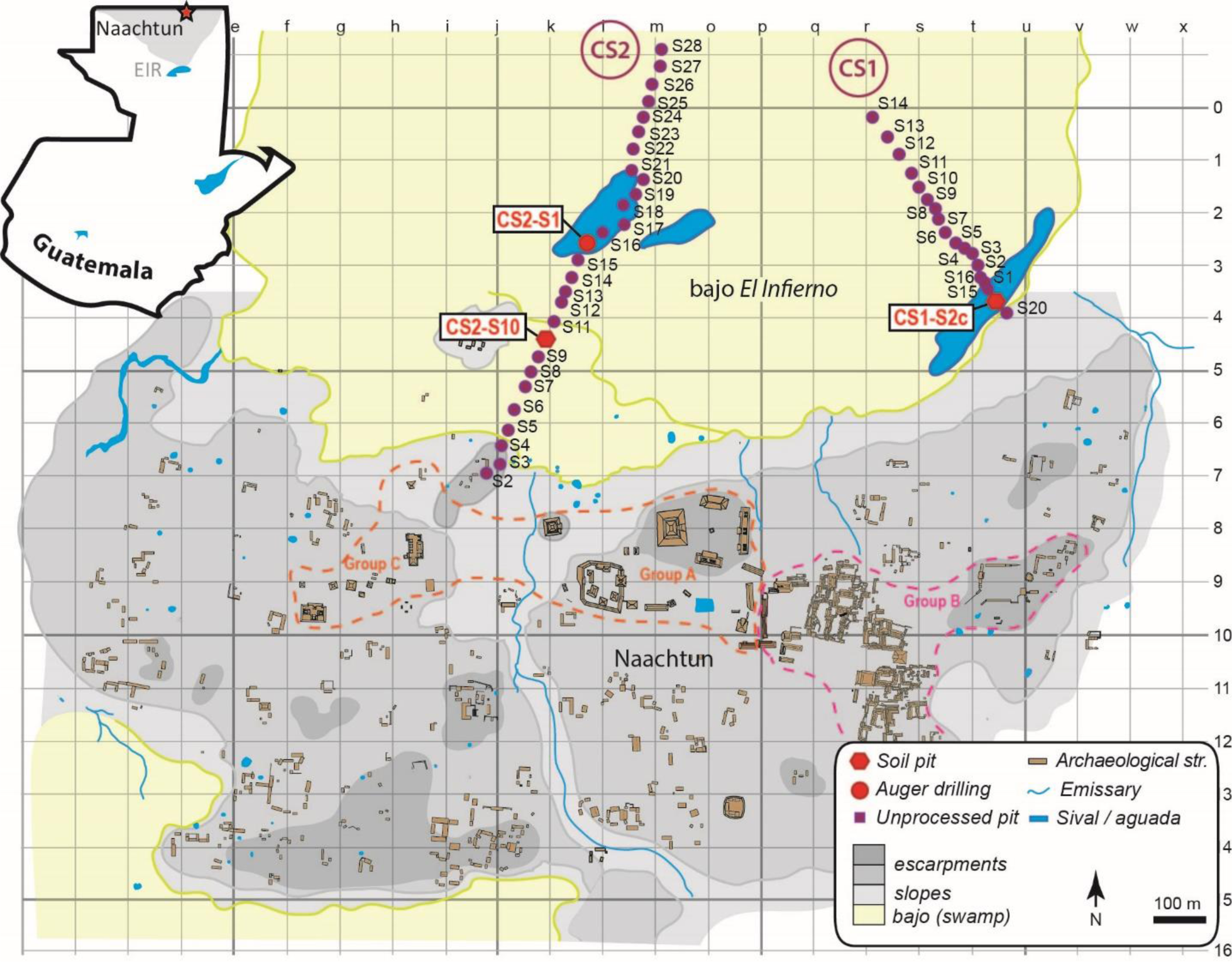
Location of the Naachtun site in Guatemala (EIR = Elevated Interior Region) and simplified site plan showing the two eastern and western sections of sedimentary boreholes.

After a scattered occupation of the site in the recent Preclassic period (400-200 BCE; Hiquet, 2020), the city of Naachtun is founded around 150 CE, at the Preclassic to Classic transition. Simultaneously, large cities in the northern Petén, such as El Mirador, Nakbe, or Ceibal, were abandoned (Wahl et al., 2007; Inomata et al., 2017), leading to population movements in the region. Located between Tikal and Calakmul, Naachtun will play a significant political role during the Classic period, notably through alliance games until the beginning of the 9th century (Nondédéo et al., 2016). From 830 CE onwards, the city gradually emptied before being definitively abandoned around 950-1000 CE (Nondédéo et al., 2013; Dussol et al., 2019; Hiquet, 2020).

### 2.2. Geographical Features of Naachtun

#### 2.2.1. Geomorphological and biogeographical contexts

The north of the Petén corresponds to the southern margins of a vast karst plateau (Elevated Interior Region; Fig.1) that extends to the north of the Yucatán Peninsula (Dunning et al., 2012). The Naachtun region presents a landscape of hills and escarpments where drainage is mainly underground, causing infiltration structures in the form of large polje-type depressions, known as *bajo* (Dunning et al., 1998). These *bajos* often take the form of marshes that are flooded during the rainy season from May to November (Hastenrath, 1976; Hidalgo et al., 2017), and which may retain residual water bodies (*Sival*) during the dry season, and more rarely, large bodies of water (*Laguna*). In contrast to the steep areas with shallow rendosols (< 50 cm), the very clayey vertisols of the *bajos* reach greater depths (1 to 2 m). The Maya indeed used these areas for hydraulic and agrarian purposes (Dunning et al., 1998 ; 2019).

Plant communities in Naachtun were first described in the 1930s by Lundell (1937) and have recently been studied again (Testé et al., 2020). The hilly areas of the site are covered with high semi-persistent forests (30-40 m) named *Ramonal*/*Zapotal* rich in trees such as Burseraceae (*Bursera simaruba*, *Protium copal*), Fabaceae (*Lonchocarpus guatemalensis*), Meliaceae (*Swietenia macrophylla*), Moraceae (*Brosimum alicastrum*), or Sapotaceae (*Manilkara zapota*, *Pouteria* sp.). The *Ramonal*’s undergrowth is relatively scattered and characterized by *xate* palms (*Chamaedorea elegans*) and *cordoncillo* shrubs (*Piper* sp.), while these species are less dominant in the undergrowth of *Zapotal*, sometimes replaced by palms of the genus *Sabal* sp. The undergrowth of the upland forests covered by dense thickets of *carrizo* bamboo (*Rhipidocladum bartlettii*) characterizes the areas of *Carrizal*.

Low forests (5-8m) cover the areas of the *bajo* (5-8 m), where plant communities are adapted to clay soils and seasonal flooding and desiccation periods (Lundell, 1937). These are mainly composed of small trees belonging to the families Anacardiaceae (*Metopium brownei*), Euphorbiaceae (*Croton guatemalensis*), Fabaceae (*Acacia* sp., *Mimosa* sp.), and Myrtaceae (*Eugenia* sp.). The structure of the *bajo* forests varies according to the areas of the depression. Near the slopes and in non-flooded areas, the undergrowth dominated by palm trees (*Cryosophila stauracantha*, *Sabal* sp.) characterizes the areas of *Escobal*. As one moves away from the slopes, palms become scarce and give way to a dense forest of small, heterogeneous, and diverse trees, referred to in our work as *Chechemal* (Testé et al., 2020). Near *Sival* areas, plant communities take the form of a monospecific lowland forest of *tinto* (*Haematoxylum campechianum*), which designates the eponymous ecosystems of *Tintal*. The residual water area, the *Sival*, is surrounded by a herbaceous plant aureole composed of Araceae (*Pistia stratiotes*), Cyperaceae (*Cyperus articulatus*), and Poaceae (*Phragmites australis*) and rare *zarza* shrubs (*Mimosa pigra*) (Lundell, 1937; Testé et al., 2020).

#### 2.2.2. Geography of Naachtun

The archaeological site of Naachtun consists of a political epicenter of 33Ha, made up of three monumental groups A, B, C, themselves distributed from west to east along two hills separated by a talweg (Fig. 1). A vast residential area of 150 Ha covers the hills around the ceremonial center with no less than 600 archaeological structures (Nondédéo et al., 2014). The LIDAR coverage highlighted that all Naachtun territory areas, apart from the *bajos*, had a density comparable to this residential area (Canuto et al., 2018).

Naachtun is bordered to the north and south by two low-lying, karst polje-type areas called *bajo* (Beach et al., 2003; Castanet et al., 2016). The deeper northern *bajo*, named *El Infierno*, is fed by several tributaries, intermittently active during rainy periods, two of which delimit the site’s residential area to the west and east. Therefore, this *bajo* is seasonally watered and has at least two perennial wetlands during the dry season, known as *Sival*, in the center and east (Fig.1) (Castanet et al., 2022). This palustrine and seasonal environment implies sedimentary, alluvial, and colluvial dynamics conducive to archiving fossil environmental bioindicators.

## 3. Sedimentary, morphostratigraphic and chrono-stratigraphic frameworks of sedimentary archives

The study of the sedimentary depositing of the *bajo* (karst polje) is based on the execution and study of 44 sedimentary boreholes implanted along two main sections, west and east (CS2 and CS1) (Fig.1; Fig.2). In addition, 15 radiocarbon datings allow a chronological framework to be constructed (Fig.2; Tab.1). The morphostratigraphy and the chronostratigraphic framework of the *El Infierno bajo* sediment deposits have thus been established (Castanet et al., 2022).

**Figure 2-.**
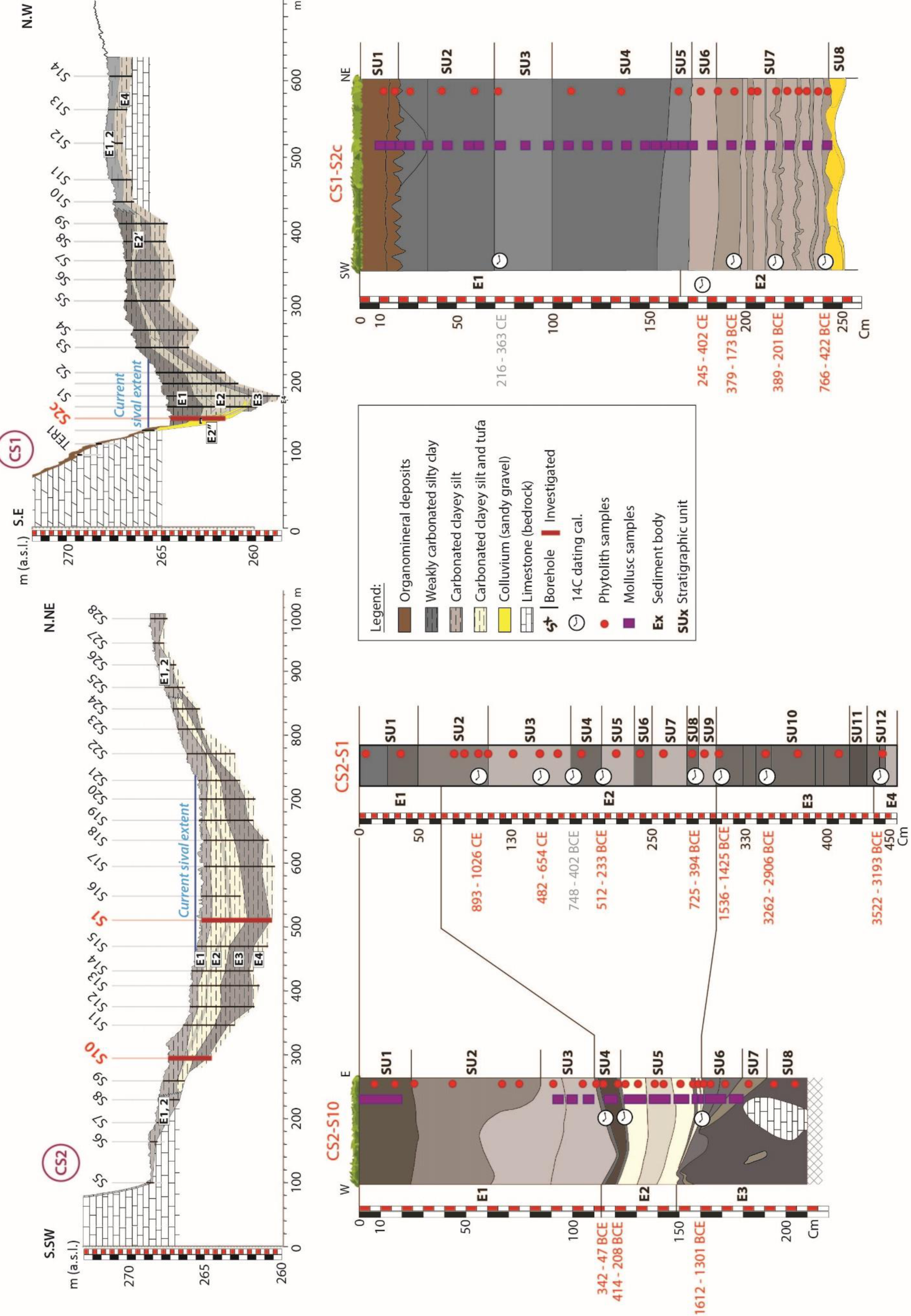
Study of the sedimentary archives of the bajo. Description of the stratigraphy and the lithology of the three sedimentary boreholes selected sampling positions and radiocarbon datings.

**Table 1-.** Radiocarbon date description table *Radiocarbon ages were calibrated using the OxCal v4.4.2 (Ramsey 2009) according to the atmospheric data from IntCal20 (Reimer et al. 2020).

| Sedimentary series | Depth (cm) or SU | Lab. ref | Radiocarbon date (BP) | Material | Calibration two sigma range (cal. BCE/CE) |
| --- | --- | --- | --- | --- | --- |
| CS2_S1 | 101-101,5 | Poz-68787 | 1070 ± 30 BP | Organic fine alluviums | [cal CE 893 : cal CE 929] 23,6 %<br>[cal CE 944 : cal CE 1026] 71,8 % |
|  | 154-157 | Poz-68448 | 1480 ± 40 BP | Organic fine alluviums | [cal CE 482 : cal CE 492] 1 %<br>[cal CE 536 : cal CE 654] 94,5 % |
|  | 182-183 | Beta-474443 | 2420 ± 30 BP | Organic fine alluviums | [cal BCE 748 : cal BCE 688] 13,9 %<br>[cal BCE 666 : cal BCE 642] 5,9 %<br>[cal BCE 567 : cal BCE 402] 75,6 % |
|  | 204-205 | Beta-474444 | 2330 ± 30 BP | Organic fine alluviums | [cal BCE 512 : cal BCE 508] 0,2 %<br>[cal BCE 481 : cal BCE 358] 91,4 %<br>[cal BCE 278 : cal BCE 258] 2,4 %<br>[cal BCE 246 : cal BCE 233] 1,4 % |
|  | 280-281 | Beta-474445 | 2390 ± 30 BP | Organic fine alluviums | [cal BCE 725 : cal BCE 706] 3 %<br>[cal BCE 664 : cal BCE 651] 2,1 %<br>[cal BCE 545 : cal BCE 394] 90,4 % |
|  | 307-309 | Poz-68450 | 3225 ± 30 BP | Organic fine alluviums | [cal BCE 1536 : cal BCE 1425] 95,4 % |
|  | 348-349 | Poz-68728 | 4385 ± 35 BP | Organic fine alluviums | [cal BCE 3262 : cal BCE 3251] 1 %<br>[cal BCE 3100 : cal BCE 2906] 94,5 % |
|  | 448-449 | Poz-68729 | 4620 ± 40 BP | Organic fine alluviums | [cal BCE 3522 : cal BCE 3336] 93,7 %<br>[cal BCE 3211 : cal BCE 3193] 1,7 % |
| CS2_S10 | 110-111 | Poz-75416 | 2115 ± 30 BP | Organic fine alluviums | [cal BCE 342 : cal BCE 323] 4,4 %<br>[cal BCE 201 : cal BCE 47] 91,1 % |
|  | 121-122 | Poz-75251 | 2305 ± 35 BP | Organic fine alluviums | [cal BCE 414 : cal BCE 351] 66,4 %<br>[cal BCE 302 : cal BCE 208] 29,1 % |
|  | 160-161 | Poz-75252 | 3190 ± 60 BP | Organic fine alluviums | [cal BCE 1612 : cal BCE 1374] 87,9 %<br>[cal BCE 1351 : cal BCE 1301] 7,6 % |
| CS1-S2C | 71-73 | Poz-68786 | 1775 ± 30 BP | Organic fine alluviums | [cal CE 216 : cal CE 363] 95,4 % |
|  | SU-6 | Poz-90901 | 1740 ± 30 BP | Charcoal | [cal CE 245 : cal CE 402] 95,4 % |
|  | 185-186,5 | Poz-70741 | 2205 ± 35 BP | Organic fine alluviums | [cal BCE 379 : cal BCE 173] 95,4 % |
|  | 227-229 | Poz-68451 | 2235 ± 30 BP | Organic fine alluviums | [cal BCE 389 : cal BCE 342] 24,2 %<br>[cal BCE 321 : cal BCE 201] 71,2 % |
|  | 245-246 | Poz-68727 | 2470 ± 30 BP | Organic fine alluviums | [cal BCE 766 : cal BCE 465] 93,4 %<br>[cal BCE 436 : cal BCE 422] 2,1 % |

### 3.1. West section (CS2)

Section CS2, located west of the *bajo*, reveals fine alluvial and colluvio-alluvial sediment deposits. 4 main sediment bodies (SB) are identified (CS2-E4 to CS2-E1). They present a tabular geometry of sediment basin filling (Fig.2). Detrital weakly carbonated silty clay deposits, poor in organic matter, alternate with carbonated clayey silt deposits with low organic matter content.

#### 3.1.1. Auger drilling CS2.S1

The CS2.S1 drilling is located in the current *Sival* in the center of this small basin. The deposits there exceed 4.5 m thick (Fig.2). It comprises 12 stratigraphic units (SU1 to SU12) grouped into three main SB (E3 to E1). Radiocarbon datings reveal the age of the deposits, including the last 5000 years (Fig.2; Tab.1).

The basal SB studied here, CS2-E3, has 1.50 meters thick (440 to 308 cm). It is composed of silty clay deposits with low carbonate content. The age of the roof of this SB is around 1536-1425 BCE (Fig.2). The median SB CS2-E2 (308 to 70 cm) corresponds to more carbonated clayey silt deposits. These deposits were established between ∼1475 BCE et ∼1250 CE according to the age-depth model developed by Castanet et al. (2022). Finally, the upper SB, CS2-E1 (CS2.S1-SU2 and CS2.S1-SU1), constitutes the upper part of this sedimentary series (Fig.2). It consists of weakly carbonated silty clay. Its base is dated around ∼1250 CE.

#### 3.1.2. Borehole pit CS2.S10

Survey (borehole pit) CS2.S10 was carried out halfway between the current *Sival* and the bottom of the *bajo* slope. It is in a currently dewatered zone for most of the time. The thickness of the deposits exceeds 2 m. The series consists of eight SU (CS2.S10-SU8 to CS2.S10-SU1) equally divided into three main SB, CS2-E3 to CS2-E1. The three SB (E3 to E1) of CS2.S1 and CS2.S10 are correlated in the CS2 reconstructed stratigraphic section model (Fig.2).

The CS2-E3 basal SB (210 to 160 cm) comprises 50 cm of weakly carbonated silty clay alluvial deposits. The median package CS2-E2 (160 to 112 cm), 48 cm thick, corresponds to a carbonated clayey silt and carbonate tufa deposit, rich in gastropod shells. Rare millimeter-scale clayey levels of detrital origin intersperse this level. Pedogenized alluvial deposits corresponding to CS2.S10-SU4 (10 cm thick), interpreted as a paleosol, constitute its uppermost part (roof). The base of CS2.E3 is posterior to 1615-1303 BCE, while its roof is dated around 175 BCE (Fig.2). The upper part of the series, CS2-E1 (112 to 0 cm), is weakly carbonated silty clay.

The geometry of the SU observed in stratigraphy reveals significant post-depositional deformations of the original sedimentary structures. The SU of the upper SB seems to be the most impacted. The processes at the origin of these deformations could be those at the origin of argilliturbation, as Beach (2009) described.

### 3.2. East section CS1 – the borehole pit CS1.S2c

The sedimentary depositing of this eastern part of the E*l Infierno bajo* is relatively similar to the western section (Fig.2). The main differences are the location of the current *Sival* immediately at the bottom of the *bajo* slope, the presence of coarser colluvium intercalated in the alluvium, and the increased thickness of the series (more than 6 m).

Survey (borehole pit) CS1.S2c was carried out at the bottom of the *bajo* slope in the *Sival*. The sedimentary series presents two large SB, CS1-E2 and CS1-E1 and 8 SU (Fig.2).

The basal SB, CS1-E2 (256 to 166 cm), has coarse colluvium at its base (CS1.S2-SU8). It then comprises a carbonated silty-clay deposit, weakly organic and stratified (CS1.S2-SU7, thick: 60cm). The dating obtained at the base of the alluvial deposits of this unit indicates an age of 768-431 BCE, while this of the upper part of this SB (CS1.S2-SU6) is established at 237-384 CE.

The upper SB, CS1-E1 (166 to 0 cm), comprises weakly carbonated silty clay to clayey silt. The SU geometry of the CS1-E1 package also reveals significant post-depositional deformation of the original sedimentary structures (argilliturbation).

## 4. Studying paleoenvironments using bioindicators

### 4.1. Study Sampling

For the three surveys of the paleoenvironmental study (CS2.S1, CS2.S10, CS1.S2C), sample selection was based on SU and radiocarbon dates. For the study of phytoliths, at least one-centimeter thick sample was taken for each SU. For the SU thick (more than 50 cm) or of sedimentary and/or pedological interest, several samples were taken. A total of 63 samples of 1 cm thickness, for 5 to 10 cm of diameter, were taken from the three profiles for the study of phytoliths (Fig.2).

Mollusks were studied on the sedimentary series CS2.S10 and CS1.S2c. Auger drilling sampling of CS2.S1 did not provide sufficient sediment volume for this type of study (Fig.2). In borehole CS2.S10, sampling was concentrated on the carbonate ensemble CS2-E2 because of its richness in gastropod shells, but also on the transition zones with the lower and upper ensembles. 11 sedimentary samples of 0.5 L to 4.5 L were collected in thicknesses of 4 to 12 cm, SU by SU, or sub-SU by sub-SU. In borehole CS1.S2c, 36 malacological samples of 5 to 10 liters were collected in the stratigraphic column for the CS1-E1 Package. Package CS1-E2 was collected as six contiguous drilling augers from the stratigraphic sets, for a total of 15 samples of 0.6 liters.

### 4.2. Application of phytoliths

#### 4.2.1. Morpho-ecological classification

Due to the production of many common phytoliths between plants, taxonomic biases usually lead to difficulties in linking a morphotype to a taxon more inclusive than the family (Rovner, 1971). This multiplicity that morphotypes are no longer associated with a taxon by name but with a parallel nomenclature based on their morphology (Madella et al., 2005; ICPT, 2019). In this article, we have chosen to follow the rules laid down by the latest published Phytolith nomenclature (ICPT, 2019) to name the relevant morphotypes. Table 2 lists the selected morphotypes and their equivalents published in Testé et al. (2020).

**Table 2-.**
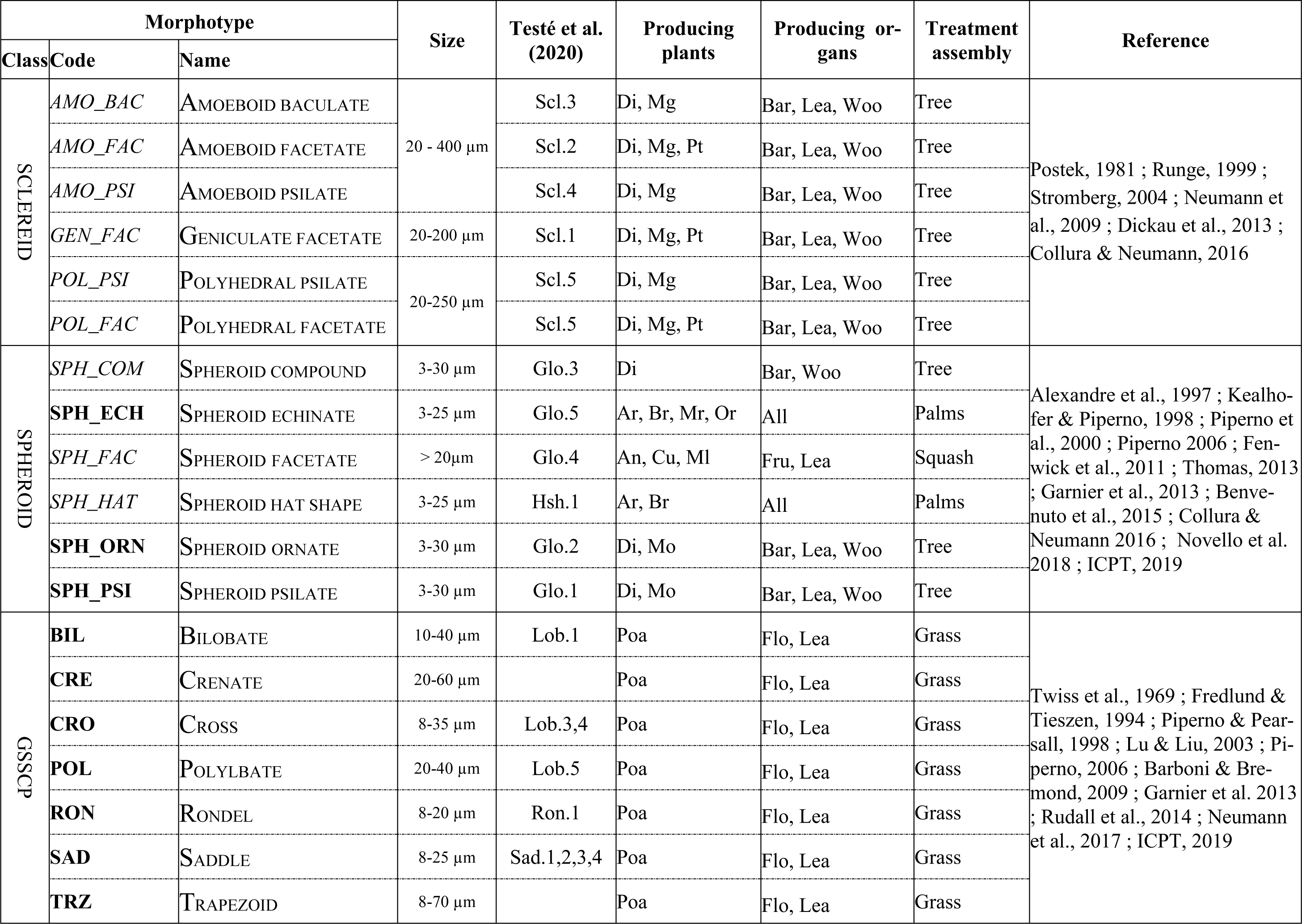
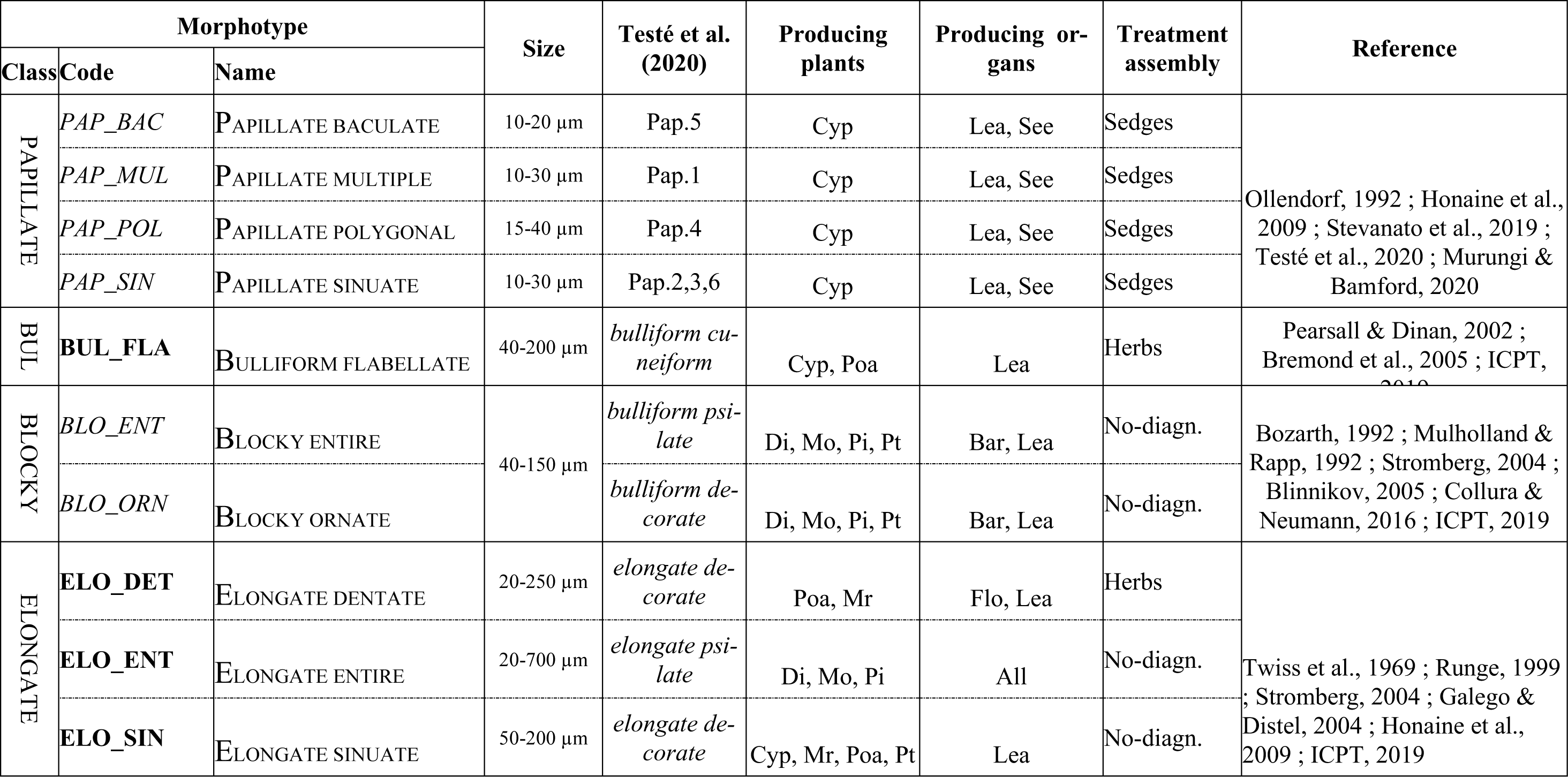
Classification of phytolith morphotypes concerning their botanical producer and an indication of treatment in assemblages. The table includes only morphotypes that run into assemblages. Morphotype codes accepted in the 2.0 nomenclature are shown in bold, those not accepted are shown in italics. Abreviated taxa : An - Annonaceae, Ar - Arecaceae, Br – Bromeliaceae, Cu – Cucurbitaceae, Cyp – Cyperaceae, Di = Dicots, Mg – Magnoliaceae, Ml – Malvaceae, Mo – Monocots, Mr – Marantaceae, Or – Orchidaceae, Pi – Pinophyta, Poa – Poaceae, Pt – Pteridophyta. Abreviated organs : Bar – bark, Flo – flower, Fru – fruits, Lea – leaves, See – seeds, Woo – wood.

AMOEBOID, GENICULATE or POLYHEDRAL morphotypes, ornamented (BACULATE, FACETATE) or PSILATE, are classified as SCLEREID class (Tab.2) and indicate mostly the presence of tree and woody angiosperms (Dicotyledons, Magnoliideae - Postek, 1981; Runge, 1999; Strömberg, 2004).

The SPHEROID class groups the sphere-like body together (Tab.2). The ornamentation of the morphotypes has been considered as an ecological type distinctiveness of the plant. The smooth, composed and granular shapes, respectively SPHEROID PSILATE, SHEROID COMPOUND and SPHEROID ORNATE, are interpreted as phytoliths of woody dicotyledons (Alexandre et al., 1997; Collura & Neumann, 2016). In the underneath, these three morphotypes together will be named as SPHEROID *mixed*. The faceted forms, SPHEROID FACETATE, are instead associated with squash and Cucurbitaceae (Piperno et al., 2000). Finally, the spiny, spherical morphotypes, SPHEROID ECHINATE, or hemispherical, SPHEROID HAT SHAPE, are associated with the Arecaceae, which produce them in large quantities (Fenwick et al., 2011; Benvenuto et al., 2015).

The small (< 25 µm), opposite-faced shapes are called GSSCP for Grass Silica Short Cell Phytolith and are produced in most subfamilies of Poaceae (Tab.2; Twiss et al., 1969; Neumann et al., 2017). Lobate shapes are found, i.e., with 2 to 5 or 6 lobes, arranged or not along a stem. Most lobates indicate the presence of Panicoidae grasses. The SADDLE morphotype has faced opposite sides, convex at the ends, and concave at the edges, giving a saddle-like appearance to these morphotypes. They are including formed in the subfamily Chloridoidae. Finally, the RONDEL has a generally cylindrical shape and where the upper face can be diagnostically ornamented as in maize, which has a wave shape (Pearsall et al., 2003), or bamboos with a spiny head (Tab.2; Piperno & Pearsall, 1998; Pearsall et al., 2003).

The many conical shapes with tabular edges form the morphotype PAPILLATE. The morphology of the cone, the number of cones, the ornamentation, and the table’s shape have many character states and make it difficult to establish a fine classification of PAPILLATE. Cyperaceae produce most of these shapes (Ollendorf et al., 1992; Honaine et al., 2009 ; Murungi & Bamford, 2020). However, in Naachtun, a particular shape PAPILLATE POLYGONAL with a polygonal and ornamented table and an angular cone, characteristic of wetland sedges (Tab.2; Testé et al., 2020), is distinguished from other morphologies (PAPILLATE SINUATE, PAPILLATE BACCULATE, PAPILLATE MULTIPLE). In the continuation of the article, these last three shapes will be treated and regrouped are the name of PAPILLATE *mixed*.

The fan-shaped morphologies give their name to the BULLIFORM FLABELLATE morphotype (Tab.2). Well marked by a somewhat curved body ending in a constricted stem, these phytoliths appear primarily in the leaves of monocotyledonous grasses (Piperno, 2006; Gu et al., 2013).

Finally, the rectangular to elongated polygonal morphologies are named BLOCKY and ELONGATE, respectively (Tab.2). These forms can have quite a diverse ornamentation or processes. However, they are present in most terrestrial plants and provide little taxonomic information. They are, however, included in counts as part of an ecological modeling approach using indices (ex: morphotype ratios).

#### 4.2.2. Assemblages, indices and statistics

##### 4.2.2.1. The general approach

The differential production of phytoliths by plants and the low taxonomic resolution lead to a general assemblage approach (Neumann et al., 2009). This approach counts all morphotypes observed in a slide until the statistical threshold of 200 diagnostic morphotypes is reached (Strömberg, 2009; Zurro, 2018). This general count is interpreted as an ecological signal because it is inherited from the production and decomposition of all local vegetation. The results obtained take the form of a diagram of relative concentrations of diagnostic morphotypes (SCLEREID, SHEROID, GSSCP and PAPILLATE, Tab.2) and ratios of non-diagnostic morphotypes (BULLIFORM, BLOCKY, ELONGATE; Tab.2) to total silica particles counted.

Within the general method, it is also possible to look for rare morphotypes characteristic of a species or genus. In our case, diagnostic phytoliths of squash and maize can be identified (Tab.2; Piperno, 2006). These identifications are relatively important since these crops are among the Maya’s agricultural heritage.

##### 4.2.2.2. Morphometric approach on CROSS morphotypes

In the absence of diagnostic maize phytoliths (Tab.2; Pearsall et al., 2003), a morphometric treatment can be applied to the shapes of CROSS, which are produced in large quantities by the maize plants attested to its presence. Pearsall and Piperno (1990) proposed a method to identify maize phytoliths from the size of the CROSS shape. It is based on the identification of four size classes: SL < 11µm, ML 11< < 16 µm, L 16< < 20 µm and XL > 21 µm. They pointed out that maize produced higher proportions of L-class CROSS than most other herbaceous plants and that it was the sole producer of CROSS XL. Iriarte (2003), who used this method, showed that no plant in the Uruguayan savannahs produced such high L morphs, or simply XL morphs.

We, therefore, chose to test this method for samples rich in GSSCP and especially in CROSS morphotypes (>10%).

##### 4.2.2.3. Numerical Approach: Index and Statistics

The counts obtained by the general method make it possible to develop numerical approaches: indices to model ecological states and statistical processes to validate diagram observations.

An index creates numerical ratios of morphotypes from counts, whose value ranges could reflect an ecological signal. In forested areas, the D/P index estimates vegetation cover from the ratio of phytoliths produced by woody dicotyledons to those produced by Poaceae (Alexandre et al., 1997). Values less than 1 indicate areas with relatively open vegetation, while values greater than 1 indicate wooded areas. Since its description, the formula has been adapted to different ecosystems (Bremond et al., 2005; Bremond et al., 2008; Garnier et al., 2013). In this study, we use the formula calibrated to the forest environments of the Petén (Testé et al., 2020), whose formula is :

D/P = (SCLEREID + SPHEROID *mixed*) / (GSSCP + PAPILLATE)

The LU index, developed from modern assemblages of Naachtun phytoliths, seeks to identify mixed environments of forest and herbaceous areas (Testé et al., 2020). This index expresses according to the formula :

LU = (Sclereid + Spheroid *mixed* + GSSCP + Papillate) / ((Sclereid + Spheroid *mixed*)*(GSSCP + Papillate))

It suggests the presence of a mixed or transition zone between the tree and herbaceous formations if its value is less than 0.25. Conversely, the higher a value is above 0.25, the more it seems to indicate a plant community dominated by one stratum. Its application in this study makes it possible to test this index to fossil assemblages and to see if it can identify transition phases in past ecosystems.

A large number of samples (63) and the statistical threshold (200 morphotypes counted) allow us to treat the fossil assemblages statistically. This latter links a cluster of similarities (Hierarchical Cluster Analysis) to a Correspondence Analysis. The HCA allows us to check whether the groupings made using the general approach have a statistical reality. The identified clusters thus serve as a control group for interpreting the CA.

The CA consist of two phases: the first corresponds to the analysis of fossil data, while the second phase integrates modern assemblages of clearly identified environments (Testé et al., 2020). This methodology allows us to verify whether the assemblages obtained by the HCA can be interpreted ecologically and whether they are similar to the modern environments of Naachtun.

#### 4.2.3. Modern reference and uniformitarianism in Naachtun

The work to characterize Naachtun’s ecosystems using phytoliths rests upon constructing a modern reference frame of assemblages about the plant communities (Testé et al., 2020). This study consists of vegetation surveys and phytolith sampling of surface soils in 43 ecological quadrats. It demonstrated that phytolith assemblages could discriminate between the different ecosystems of Naachtun.

The areas of hill forests (*Ramonal/Zapotal*) corresponding to the archaeological site present phytolith assemblages dominated on average by 65% SPHEROID *mixed* (as a reminder: the PSILATE, ORNATE and COMPOUND shapes) and 18% SPHEROID HAT SHAPE morphotypes, respectively produced by woody dicots trees and certain families of Arecaceae. The bamboo thicket areas (*Carrizal*) distributed in hill forests are dominated by 60% SPHEROID *mixed* and 20% RONDEL *crested* morphotype identified in many bamboo species.

A forest rich in palm trees (*Escobal*) covers the slopes and bottoms of hill environments. Phytolith assemblages in these environments are 50% dominated by SPHEROID *mixed* and 40% by SPHEROID ECHINATE, produced by certain families of Arecaceae.

Plant communities in the *bajo* evolve from mixed lowland forests (*Chechemal*) on the margins to monospecific lowland forests (*Tintal*) around perennial water areas that are palustrine environments dominated by grasses and sedges (*Sival*). *Chechemal* phytolith assemblages are dominated on average 80% by SPHEROID *mixed* and have residual sponge spicules around 10%. *Tintal* areas have more diversified assemblages with an average of 40% SPHEROID ECHINATE, 20% GSSCP produced by the Poaceae, and 20% PAPILLATE produced by the Cyperaceae. The *Tintal* areas constitute a transition zone with the *Sival* environments whose assemblages are 15% SPHEROID *mixed*, 40% GSSCP, and 40% PAPILLATE. The assemblages of the *Tintal* and *Sival* zones are composed on average of 30% sponge spicules and 5 to 10% diatoms.

It is based on these modern assemblages, calibrated on the modern ecosystems of Naachtun, that the fossil assemblages studied in the *bajo* of Naachtun are interpreted. However, this interpretative basis does not allow the study of anthropogenic or currently foreign environments in Naachtun. Also, these results show the need to cross-reference the phytolith data with complementary bioindicators.

### 4.3. Multiproxy study: crossover of bioindicators

#### 4.3.1. Spicules and Frustules

The modern study also demonstrated the interest of taking into account other siliceous microfossils preserved in these deposits, such as sponge spicules and diatoms. They complement the paleoecological information, particularly on the hydrological state of the ecosystems.

These siliceous microfossils are observed from phytolith slides and are therefore included in particle counts. However, due to the lack of regional classification, their treatment is not taxonomic but numerical. Only the concentrations of these two types of bioindicators are calculated based on the total number of particles counted. This accounting approach made it possible to identify the water availability of the quadrats for the modern reference (Testé et al., 2020).

#### 4.3.2. Modern and fossil mollusks

##### 4.3.2.1. Preparation and study of malacological samples

After drying, all samples were sieved with Sodium Hexametaphosphate at 40g/L on a 500 µm mesh. The sieve rejects were sorted into two fractions : a detrital fraction comprising minerals, organic and inorganic residues including portions of non-identifiable shells, and a fraction composed solely of identifiable shells. Counting consists of identifying, for each species, the minimum number of individuals by adding to the whole shells the avered jumble of apexes and openings. A sample is statistically representative when it exceeds 200 individuals (Evans, 1972). All counts are then plotted in a distribution diagram according to stratigraphic records. Naachtun sediments are also rich in fossil oogons of Characeae. We decided to evaluate their relative richness in our malacological samples because of their ability to sign water conditions in stagnant water bodies (Soulié-Märsche, 2002).

##### 4.3.2.1. Modern fauna of gastropods

The taxonomic determination of fossils only depends on conchological characters. A taxonomic and ecological survey of terrestrial and freshwater mollusks was led in 1937 by Goodrich and Van der Schalie in the Petén region. Further missions to the CMLs improved species lists for the region (Basch, 1959; Dourson and Caldwell, 2018). The taxonomy used in this work follows the list established by Thompson (2011) on Mexico and Central America.

Among the species common to the *aguadas*, marshes, *bajos*, and *Sival* zones noted by Goodrich and Van der Schalie (1937) and Basch (1959) in Tikal, five shapes are in the majority in the malacological assemblages collected at Naachtun. They all correspond to species of aquatic gastropods.

The first shape found corresponds to the species *Pomacea flagellata* (Plt.1n-p), a large ampullar of up to 5 cm in height, and with significant morphological variability (Goodrich and Van der Schalie, 1937). It is an amphibious organism living both in the water body and on its margin (Naranjo-Garcia and Olivera Carrasco, 2014) and whose shells are scattered on the floor of the *bajo* (Basch, 1959). It is often found in the archaeological contexts of the Petén for culinary reasons (Moholy-Nagy, 1978). On the other hand, fossil shells are rarely preserved whole in the sediments due to their large size.

The four other shapes are all millimeter shells.

**Plate 1.-.**
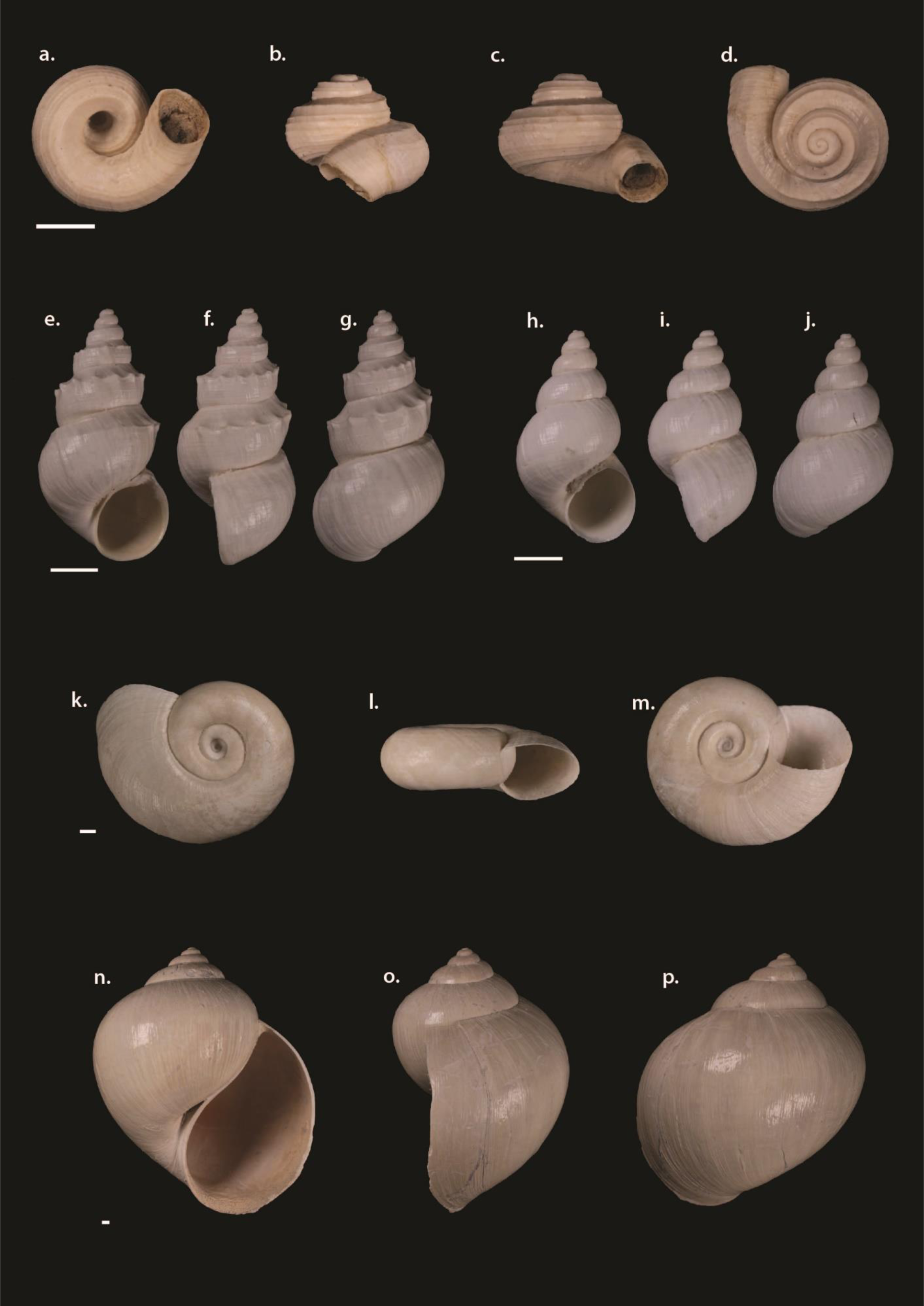
Photograph under different views of the main fossil shells found in the sediments of Naachtun. a-d = Cochliopina infundibulum; e-g = Pyrgophorus coronatus spiny-shape; h-j = Pyrgophorus coronatus smooth-shape; k-m = Planorbidae; n-p = Pomacea flagellata. Scale bar = 1 mm; (Photography by Pierre Lozouet).

Among these is the species *Cochliopina infundibulum* (Plt.1a-d). The shell is particularly recognizable by its flattened shape and multi-carina ornamentation. The species is common as a sub-fossil organism but rarer in the present (Goodrich and Van der Schalie, 1937; Covich, 1976; Kim and Rejmánková, 2002). The work on ecological indicators of riparian areas in Tabasco by Loreto (2011) considers *C. infundibulum* as a typical species of marsh landscapes.

Two other shapes can be attached to the same family of Cochliopideae. The most remarkable is a conical shell of millimeters in size with a spiny carina. In the lowland literature, it is generally assigned to the species *Pyrgophorus coronatus* (Plt.1e-g). However, recent work by Grego et al. (2019) describes a modern, more minor, pyramidal shape of the Yucatán with a spiny carina as *Pyrgophorus thompsoni*. Juveniles of *P. coronatus* already have spiny shells and make it difficult to identify possible forms of *P. thompsoni* in our counts. We observe a very significant variability in the expression of the carinas, with some shells having only rare spines on the whorl, or even the total absence of carinas in these two species of *Pyrgophorus* sp. (Covich, 1976; Grego et al., 2019). This type of variation in ornamentation appears in other *Pyrgophorus* sp. in Latin America (Nava et al., 2010). This variability is related to changes in environmental conditions such as salinity, presence of predators, availability of dissolved oxygen (Covich, 1976; Vermeij and Covich, 1978; Bradbury et al., 1990). It is difficult to divide these shells into different species on conchological characteristics. Thus in this work, these types of shells are counted as *Pyrgophorus* sp.. However, the supposed ecological interest of the variations in ornamentation in this genus leads us to count the smooth (Plt.1h-j) and spiny (Plt.1e-g) forms differently.

The final shape corresponds to flat spiral shells of the family Planorbidae. Most of the shapes identified correspond to juveniles, and the assignment to adult shapes is complicated. Among the species present in the Petén that could correspond to adult forms is *Biomphalaria havanensis*, *Biomphalaria obstructa*, *Biomphalaria petenensis*, *Planorbella trivolis* (Plt.1k-m). Because of the uncertainties related to the available samples, these shapes are counted as Planorbidae. They are freshwater organisms common in all aquatic environments (Goddrich and Van der Schalie, 1937).

As these five shapes represent almost all the assemblages, the rare shells belonging to other species have limited occurrences and will be treated as “other species” in our counts.

## 5. Past ecosystems studied by phytoliths and siliceous bioindicators

### 5.1. Study of Auger drilling CS2.S1

The CS2.S1 survey consists of 20 phytolith samples (Fig.3).

**Figure 3-.**
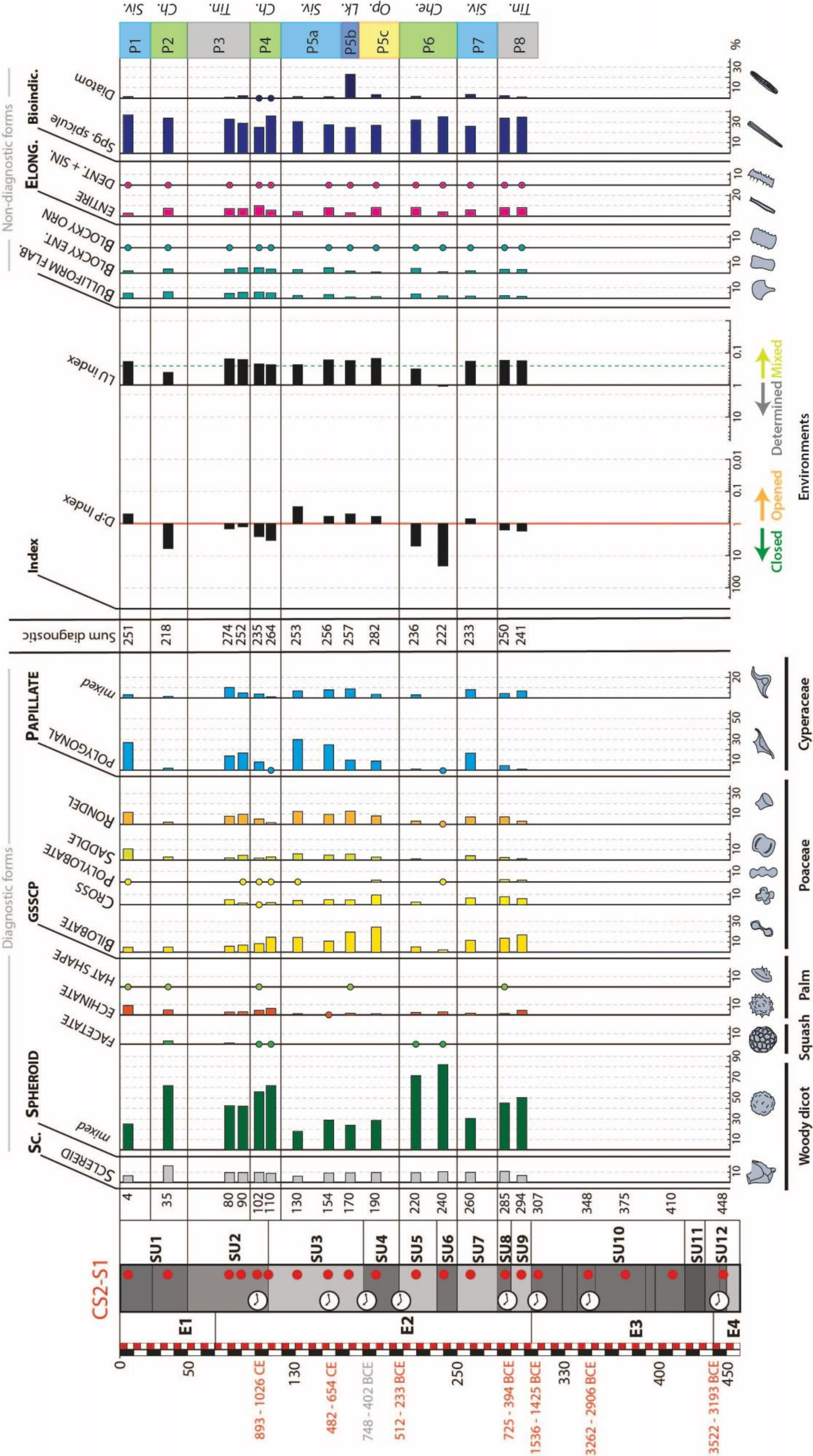
Distribution diagram of diagnostic, non-diagnostic phytolith morphotypes and siliceous microfossils from auger drilling CS2.S1. Identification of the temporal evolution of ecosystems. Red dots = samples taken from the auger drilling for phytolith studies. Che. = Chechemal; Lk. = lake; Op. = open vegetation; Tin. = Tintal; Siv = Sival.

The CS2-E3 package (CS2.S1-SU12 to SU10) is devoid of phytolites and other siliceous bioindicators. This absence could be the result of deposition conditions or diagenesis unfavorable to the conservation of siliceous remains.

The S1-SU9 and S1-SU8 at the base of the CS2-E2 package form a first phase (S1-P8) where the phytolith assemblages are dominated by 45% to 51% SPHEROID *mixed*, showing a significant proportion of woody material in the ecosystem (Fig.3). Cumulative GSSCP values reach up to 32%, including 14 to 17% BILOBATE, while PAPILLATE do not exceed 8%. This herbaceous component of the assemblage is also supported by D/P index values slightly above 1 and LU values below the 0.25 threshold, indicating a mixed ecosystem. Finally, we note sponge spicules values above 30% and residual diatom levels, characteristic of temporary wetlands. This type of assemblage is comparable to the assemblages of Naachtun *Tintal* environments, i.e., a forest with a periodic herbaceous stratum, seasonally submerged due to its proximity to a perennial water body.

S1-SU7 presents an assemblage where the SPHEROID *mixed* rate has decreased to 30% and where the GSSCP values are similar to the previous phase (31%) (Fig.3). In this S1-P7 phase, on the other hand, the PAPILLATE reach values of 25% of the assemblage. It seems that this environment is slightly more open and moist than the S1-P8 phase. D/P index values confirm this slightly below 1, but the LU index, still below 0.25, indicates an environment with a tree-like component. Phytolith and bioindicator values have changed little in this phase, sponge spicules still exceed 25%, and diatoms increase to 5%. It is possible to interpret this assemblage as the presence of a *Sival* zone, a perennial wetland dominated by grass assemblages, at the edge of the forest or a *Tintal* zone subject to a closer wetland.

The next phase, S1-P6, covers S1-SU6 and S1-SU5 (Fig.3). The assemblage is characterized by SPHEROID *mixed* rates greater than 70%, associated with a substantial decrease in GSSCP (>12%) and PAPILLATE (>5%) values. These values indicate the presence of a dense forest ecosystem. The originality of these samples lies in the residual presence of the SPHEROID FACETATE morphotype, which attests to the presence of Cucurbitaceae. The D/P index values range from 8 to 11, while the LU index values are above the threshold of 0.25, confirming the dominance of woody species in the ecosystem. We still find sponge spicules rates between 32 and 36% and shallow diatom values (> 2%), indicating the presence of water at least part of the year. This type of assemblage is identical to the assemblages of modern *Chechemal* ecosystems, i.e., a lowland swamp forest without undergrowth and subject to short seasonal flooding.

The terminal SU of the CS2-E2 assemblage (S1-SU4 and S1-SU3, except for S1-110) form an S1-P5 phase whose variations in the assemblage show an evolving ecosystem (Fig.3). SPHEROID *mixed* values attain ranging from 24 to 29% before falling to 18% in S1-130. GSSCP values vary between 30 and 47%, with a maximum meeting for SU4 (S1-190), where BILOBATE and CROSS make up 23% and 10% of the assemblage, respectively. These high proportions of morphotypes, often associated with Panicoideae (Fredlund and Tieszen, 1994; Barboni and Bremond, 2009), have not been observed in modern samples from Naachtun (Testé et al., 2020). PAPILLATE levels increase progressively during this phase from 12% in S1-190 to reach a level of 37% in the upper part of S1-SU3 (S1-130). These assemblages show rather open to herbaceous dominated environments. This analysis is confirmed by the D/P index values, which are all below 1. The LU index values, all below 0.25, however, indicate the presence of a tree component in this open formation. Sponge spicules have values ranging from 25% to 31%, while diatoms remain below 4% except for a 23% peak at the base of SU3 (S1-170). This phase would correspond to an open ecosystem where the increase of Cyperaceae on Poaceae would indicate an increase in the humidity of the environment until resembling an ecosystem close to the modern *Sival*. This S1-P5 phase could be divided into three events, S1-P5c representing an open area with herbaceous communities not known in the current one at Naachtun, an extended wetland phase indicated by the high proportions of diatoms S1-P5b, and finally, an area of *Sival* S1-P5a.

S1-SU2 includes four samples that form 2 plant phases (Fig.3). The first phase, S1-P4 (which also includes the top part of S1-SU3 with sample S1-110) has high rates of SPHEROID *mixed* (> 56%), average values of GSSCP (between 16% and 21%), and PAPILLATE (between 2 and 11%). We also note the presence of SPHEROID FACETATE phytoliths. The D/P index values are above 2.5, indicating a closed environment, but the LU index values, slightly below 0.25, indicate a mixed environment. Sponge spicules retain high values between 25 and 36%, and diatoms are poorly represented (>1%). This assemblage is similar to S1-P6 phase interpreted like a *Chechemal* forests. So, it is possible to interpret the ecosystem of the S1-P4 phase as a low forest of *bajo*. For the next phase, S1-P3, SPHEROID *mixed* decreases to 42% of the assemblages, and SPHEROID FACETATE are absent. Conversely, GSSCP remain stable at around 21%, while PAPILLATE increase to 21% and 24% of the assemblages. This increase in the grassy component in the assemblages is correlated with a decrease in D/P index values around 1. We note stabilization of sponge spicules values (38 to 42%) and diatoms (1 to 3%). These variations in the presence of grasses, particularly Cyperaceae, suggest replacing a *Chechemal*-type *bajo* forest with a more hydrophilic *Tintal*-type forest.

The base of the S1-SU1 constitutes the S1-P2 phase, whose assemblage corresponds in all respects to the S1-P4 and S1-P6 phases (Fig.3). It consists of 60% SPHEROID *mixed*, 10% GSSCP, and 4% PAPILLATE. It presents SPHEROID FACETATE, high D/P (>5), and LU (>0.40) values and spicules up to 35% when the diatoms are residual. This type of assemblage is associated with low forests of *bajo* of the *Chechemal* type.

Finally, for the terminal part of S1-SU1 (S1-4), the corresponding S1-P1 phase presents SPHEROID *mixed* values of 25% associated with GSSCP and PAPILLATE values reaching 28% and 30%, respectively (Fig.3). While GSSCP are dominated by SADDLE and RONDEL morphotypes (22%), PAPILLATE POLYGONAL shapes dominate (27%). This type of assemblage is similar to the modern aquatic ecosystem assemblages of the *Sival* type.

### 5.2. Study of Borehole pit CS2.S10

#### 5.2.1. General study of fossil assemblages

The study of borehole CS2.S10 is composed of 23 samples (Fig.4).

**Figure 4-.**
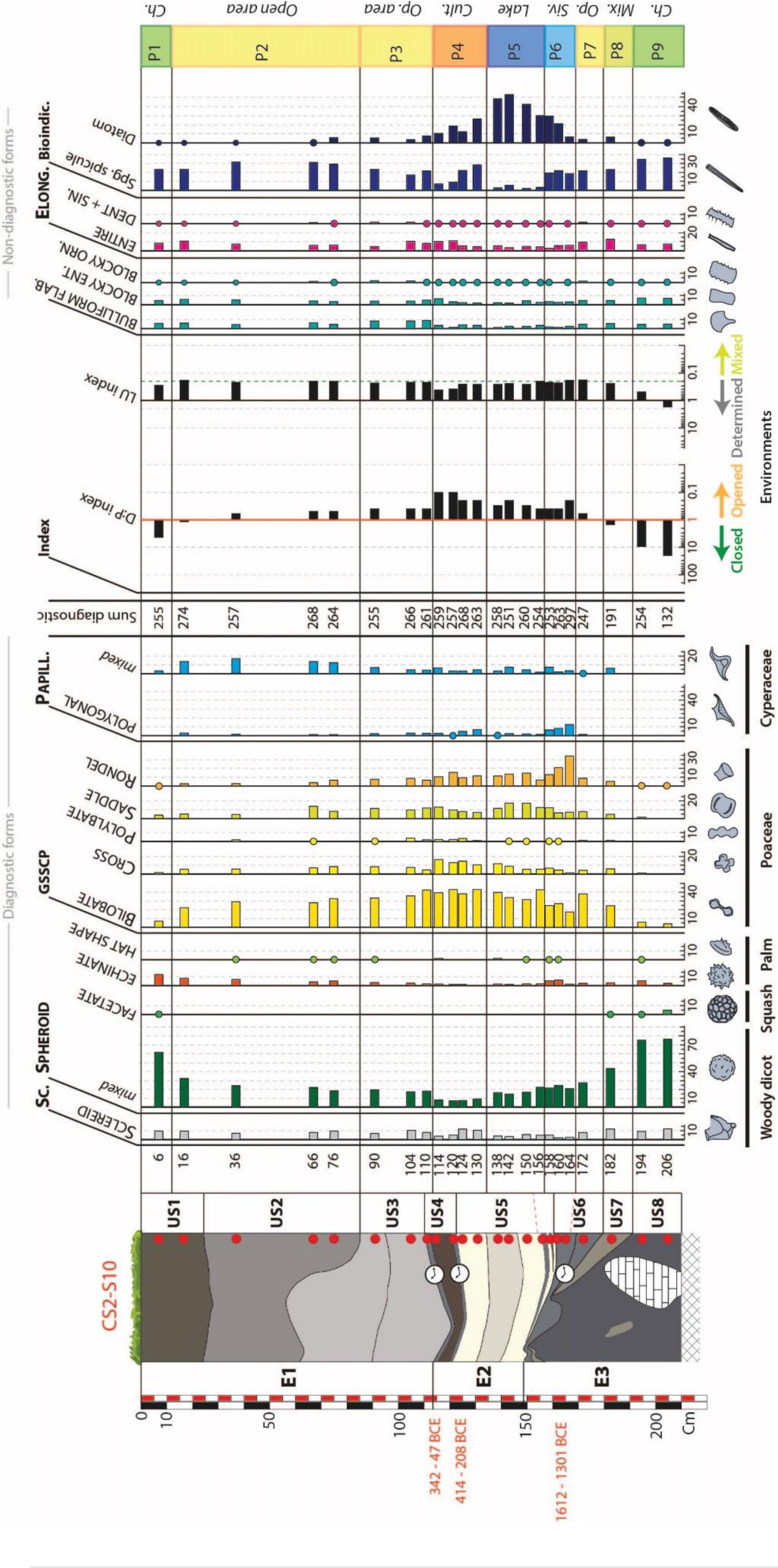
Distribution diagram of diagnostic, non-diagnostic phytolith morphotypes and siliceous microfossil from borehole pit CS2.S10. Identification of phases marking the temporal evolution of ecosystems. Red dots = samples taken from the borehole pit for phytolith studies. Che. = Chechemal; Lk. = lake; Op. = open vegetation; Tin. = Tintal; Siv = Sival.

The S10-SU8 at the base of the borehole presents two samples that correspond to the S10-P9 phase dominated by at least 75% SPHEROID *mixed* and associated with residual levels of SPHEROID FACETATE (Fig.4). GSSCP morphotypes represent less than 10% of the assemblages, and PAPILLATE are absent. D/P values are greater than 9, and LU values are greater than 0.45. Finally, among the bioindicators, we find sponge spicules in 30% of the assemblages and residual traces of diatoms. This type of assemblage is identical to the modern *Chechemal* forest assemblages.

S10-SU7 shows an S10-P8 evolution, where SPHEROID *mixed* morphotypes decrease to 42% while GSSCP reach 40%, of which 24% is BILOBATE, and PAPILLATE get 6% (Fig.4). This type of assemblage indicates the significant presence of herbaceous plants in a forest area. Thus, the D/P index decreases towards the benchmark value of 1, while the LU index drops to 0.22, indicating a mixed plant community. While sponge spicules are slightly less present with values at 22%, diatoms reach higher values around 5%. This assemblage can be interpreted as a mixed, forested area with a strong herbaceous component near an aquatic area. However, like the S1-P5 phase, high values of BILOBATE morphotypes suggest that the Poaceae communities are different from the current ones.

The base of S10-SU6 continues in this opening dynamic with the S10-P7 phase (Fig.4). SPHEROID *mixed* now represent only 28% of the assemblage, while GSSCP account for 60% of the assemblage, 38% are BILOBATE. PAPILLATE are still tiny (2%), and SPHEROID FACETATE no longer appear in the assemblage. The D/P index takes a value lower than 1 (0.6), while the LU index remains lower than 0.25. Spicules represent 26% of the particles counted, and diatoms are stable with values around 5%. This phase confirms the opening of the environment dominated by Panicoidae communities in a periodic aquatic context.

The two upper samples of S10-SU6 and the basal S10-SU5 (S10-158) sample constitute an evolutionary phase of the S10-P6 assemblages (Fig.4). The SPHEROID *mixed* remain stable during this phase with relatively low values between 20% and 23%. The GSSCP decrease softly from 60% to 56% during this phase. The RONDEL (34% to 13%) shapes became less important than the BILOBATE (17% to 25%), while the SADDLE remain relatively stable (between 7% and 12%). PAPILLATE also follow a gradual evolution from 16% to 12%. The D/P index shows values well below 1, while the LU index remains stable, with average values at 0.21. Sponge spicules remain relatively stable, with values more or less identical to the previous phase (between 22% and 26%). The gradual evolution of the phase is also well marked by the successive increase in diatom rates from 8% to 35%. The high RONDEL and PAPILLATE values during this phase correspond well to a modern *Sival* environment. However, the high diatom values and the return of phytoliths associated with Panicoidae could instead show the installation of a more perennial wetland with different grassy vegetation than the current *Sival*.

The next phase, S10-P5, is spread over the S10-SU5 up to sample S10-138. The assemblage of this phase presents relatively low SPHEROID *mixed* rates between 14% and 22% (Fig.4.). Conversely, GSSCP dominate the assemblage to 68% to 73%, of which 37% on average are BILOBATE. PAPILLATE are not very present, representing on average 5% of the assemblages. The D/P index is well below 1 (from 0.2 to 0.4) in this phase, while the LU index values are on average equal to 0.25. In this phase, the dominant plant formation corresponds to a savanna dominated by Panicoidae. A great variation occurs with a significant decrease in the rates of spicules whose values are lower than 6% and which are replaced by diatoms whose rates increase from 37% to 56%. Phytoliths here constitute a predominantly herbaceous formation, while diatoms indicate a larger size of the body of water. Thus the assemblage could correspond to a wetland with vegetation very different from modern *Sival* areas.

The top part of the S10-SU5 and the basal part of the S10-SU4 correspond to a new phase, S10-P4 (Fig.4). In these phytoliths assemblages, the SPHEROID *mixed* rates present the minimum values of the cut between 7% and 9%. Conversely, GSSCP morphotypes represent at least 71% of the assemblages, with an average of 40% for BILOBATE, while CROSS morphotypes reach values twice as high as in the other phases, ranging from 10% to 16%. PAPILLATE are always rather weakly represented with a mean value of less than 7%. D/P values are minimal, with values from 0.1 to 0.2 and LU values are slightly above 0.25, increasing to 0.4. The values of sponge and diatom spicules change during this phase. Despite reappearing, the spicules have values that increase from 21% to 9% during this phase. Diatoms follow the same phenomenon increasing from 30% to 13%. The ecosystem is dominated by herbaceous plants, including Panicoidae, whose high proportions of CROSS could suggest the presence of maize (cf. 4.2.2). The decrease in spicule and diatom rates could indicate a decrease in the water available in the environment during this phase while still indicating a perennial water phase.

However, the top sample of S10-SU4, S10-110 marks the entry into a new phase S10-P3 which also frames S10-SU3 (Fig.4.). SPHEROID *mixed* values rise from 17% to 19%, while the GSSCP decrease from 66% to 60%, BILOBATE following this trend from 41% to 34%. PAPILLATE are still relatively low even though there is a slight increase from 6% to 10%. Thus the D/P values increase but remain well below 1 (0.3-0.4) while the LU index values decrease slightly below the threshold value of 0.25 (0.21-0.22). During this phase, sponge spicules regain 25% to 30%, and diatoms decrease from 9% to 5%. The middle still corresponds to an open area at Panicoidae, but the decrease in diatom levels in favor of sponge spicules suggests that the body of water continues to decrease to a regime closer to modern *Sival*.

The S10-SU2 and the base of the S10-SU1 correspond to the S10-P2 phase, whose assemblages are evolving (Fig.4.). The SPHEROID *mixed* gradually increase from 19% to 32%. The GSSCP values decrease from 55% to 34%, still dominated by the BILOBATE (from 33% to 22%). PAPILLATE reach on average 15% of the assemblages but whose main morphotypes do not belong to PAPILLATE POLYGONAL forms but rather to PAPILLATE *mixed* shapes (Testé et al., 2020). During this phase, there seems to be a more tree-like component in an open ecosystem with a modification in grassy plant communities where Cyperaceae are present together with Panicoidae. The open character of the ecosystem is confirmed by the D/P index values, which remain below 1 (from 0.4 to 0.8), while the low tree component of the environment correlates well with the LU environment mixing index values and values well below 0.25 (from 0.20 to 0.16). The rate of spicules also decreases during this phase from 36% to 28%. At the same time, the diatoms fall below 1% of the assemblage. The variation of these bioindicators seems to confirm a dynamic of decrease of the water body already observed in the previous phases.

The last phase, S10-P1, corresponds to the upper sample of S10-SU1 (Fig.4). The SPHEROID *mixed* have a value of 62% and are again in association with the SPHEROID FACETATE. The sum of the GSSCP reaches 13%, and the PAPILLATE represent only 3% of the assemblage. The D/P index gives a value of 4, and the LU index value is 0.29. The spicules remain stable at 29%, and the diatoms represent less than 1% of the assemblage. We relate these data to a modern *bajo* forest of *Chechemal*-type.

#### 5.2.2. Morphometric distribution of CROSS

Due to the high proportions of CROSS morphotypes in CS2.S10, it was possible to study the distribution of their morphometric types (Fig.4.). Eight samples were tested on S10-SU7 to S10-SU2 with a mean count of 39 CROSS morphotypes per sample (Fig.5).

**Figure 5-.**
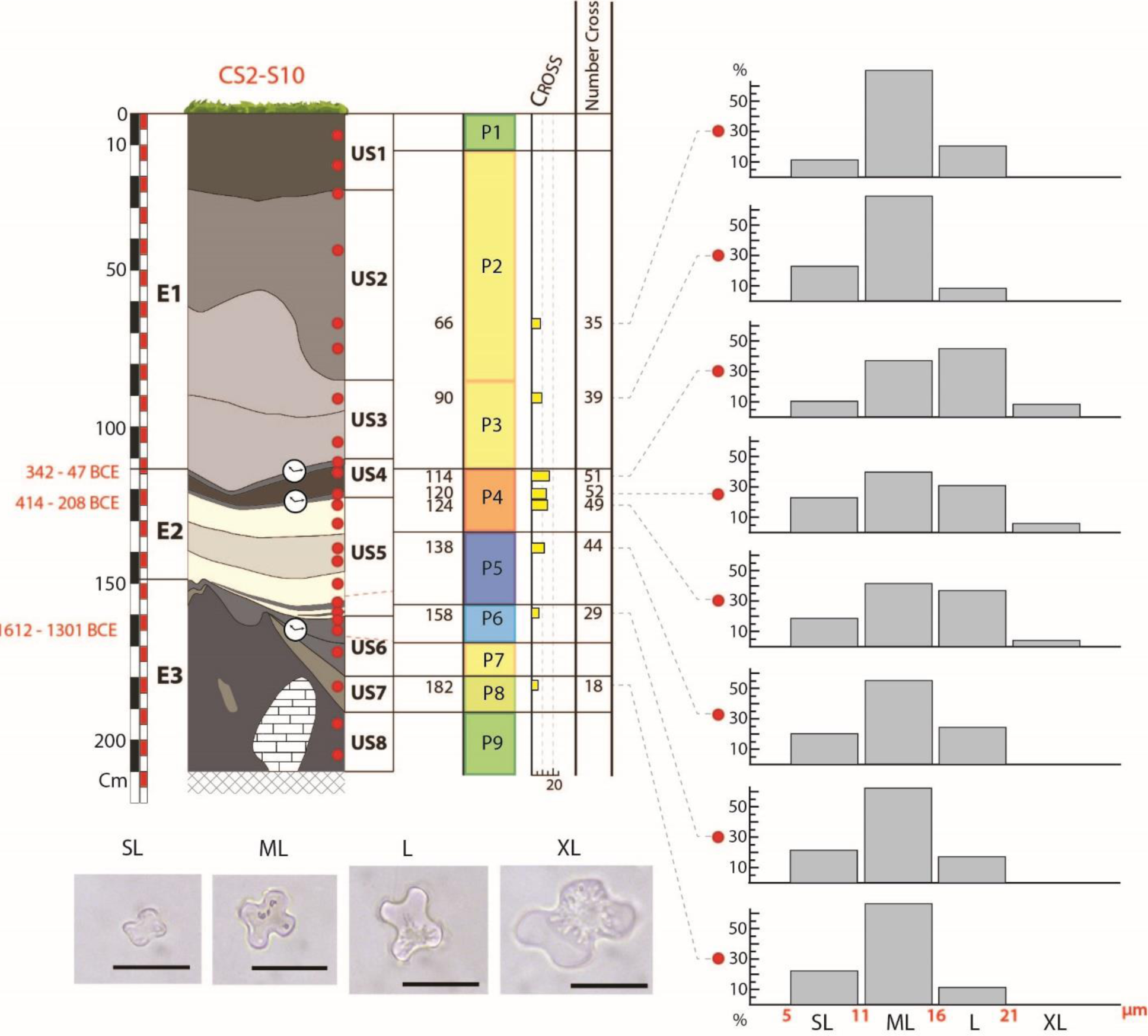
Distribution of morphometric classes of Cross morphotypes within eight samples from borehole pit CS2.S10. scale bar −20µm.. The XL morphotype, greater than 21 µm, is interpreted as an indicator of corn.

The CROSS from S10-SU7 to S10-SU5 is dominated mainly by ML size morphotypes at more than 50%. The SL morphotypes represent on average 20% of the CROSS in these units, while the L shapes are less than 20%, and the XL shapes are absent. However, in the S10-US5 sample, the values of the L and SL shapes equalize around 20%.

The upper part of S10-SU5 and basal part of S10-SU4 correspond to the S10-P4 phase, which has the highest number of CROSS in its assemblages (Fig.5). Three samples were studied there. In these, the CROSS bevy have a distribution of different morphotypes. The SL shapes are always less than 20%. The ML shapes show a less than 35% downward trend, while the L morphotypes exceed 40%. These are the only levels where a residual rate of XL shapes associated with maize is present (from 4% to 8%).

S10-SU3 and S10-SU2 are represented by two samples with maximum values for ML shapes exceeding 60%, while SL and L forms are consistently below 20% of the CROSS assemblages. The XL morphotypes are absent from these assemblages.

Based on these results (Fig.5), only the S10-P4 phase shows clear indications of maize cultivation among the Panicoidae signal. This level also corresponds to a paleosol, and the high levels of spicules and diatoms suggest that this farming takes place in a relatively humid environment (Fig.4).

### 5.3. Study of Borehole pit CS1.S2c

The phytoliths’ study in borehole pit CS1.S2c is based on 20 samples (Fig.6). The basal unit S2-SU8 was not samplied. S2-SU7 is studied from 8 basal samples forming the S2-P4 phase. The SPHEROID *mixed* represents, on average, 26% of the assemblages, with values ranging from 17% to 32%. SCLEREID has a strong presence with average values of 20%, of whom peaks at 40% on the basal sample of this phase. While SPHEROID ECHINATE show low rates compared to the western section (average 6%), this phase is the only one to show such high SPHEROID HAT SHAPE rates (up to 9%). On average, GSSCP represent 32% of the assemblages, with BILOBATE and SADDLE dominating, respectively at 15% and 9%. The PAPILLATE, dominated by MULTIPLE and SINUATE morphs, have values that vary from 15% for the basal and summit samples of the phase to 6% for the median part. However, the high values of PAPILLATE *mixed* type suggest that the Cyperaceae communities are different from those found in the modern areas of *Sival*. Based on these assemblages, the S2-P4 ecosystem could be interpreted as a forest savannah or a forest with significant herbaceous undergrowth. Because the borehole pit is located at the foot of the slope, these assemblages could also correspond to a composite record of forest vegetation from the hillside and a herbaceous component composed of Panicoidae and Cyperaceae from the *bajo* areas. These analyses are confirmed by the D/P index, which varies between 0.7 and 2.3, whose an average value of 1.1. Equally, all LU index values are below 0.25 (between 0.16 and 0.19). Sponge spicules represent on average 15% of the particles while diatoms are residual (>1%), indicating the presence of a wetland nearby, but whose action would be somewhat seasonal, leaving the ecosystem emergent for a large part of the year.

**Figure 6-.**
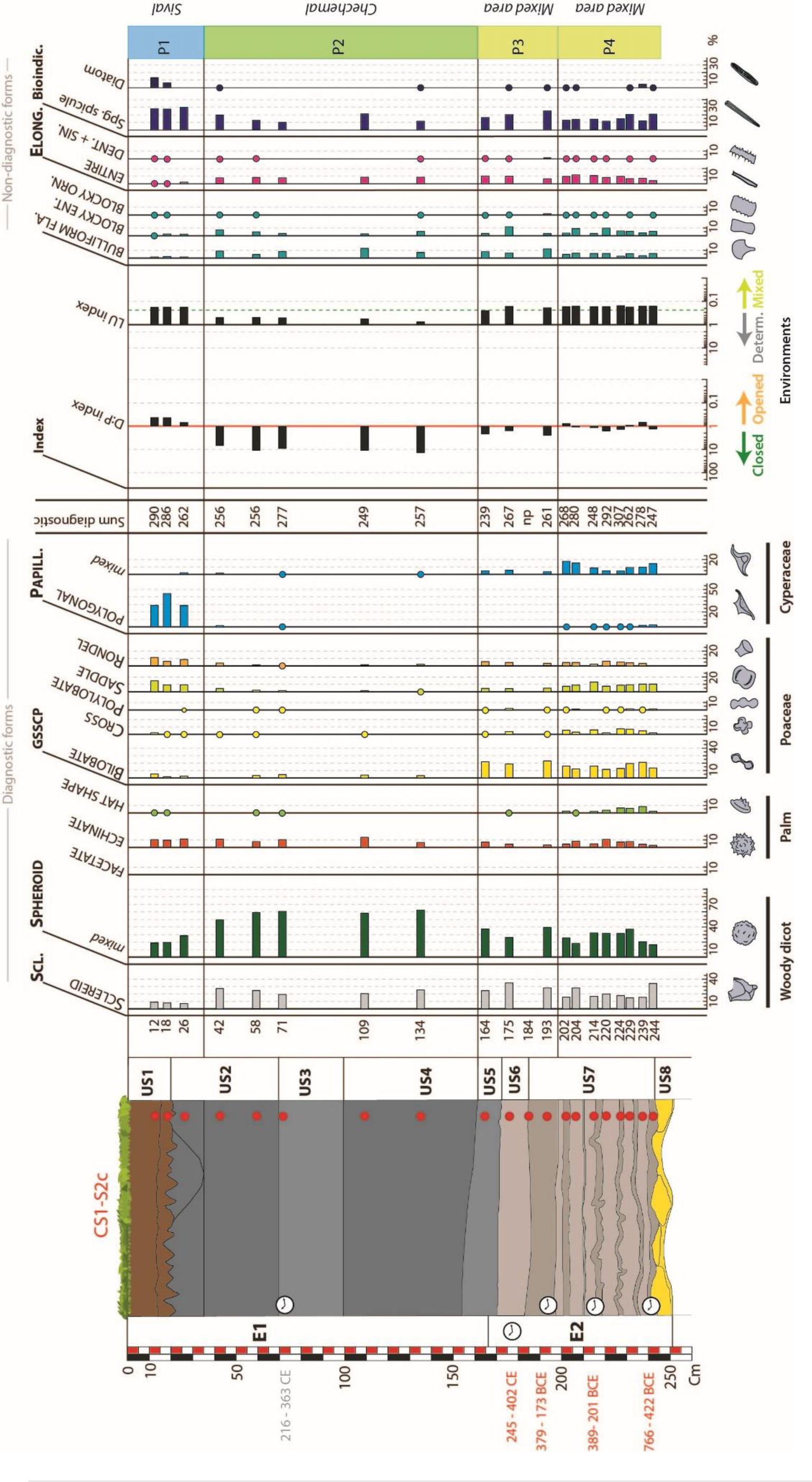
Distribution diagram of the diagnostic and non-diagnostic phytoliths shapes, and siliceous microfossils in borehole pit CS1.S2c. Identification of phases marking the temporal evolution of ecosystems. Red dots = samples for the study of phytoliths. Che = Chechemal; Ope, area = Open vegetation; Siv = Sival

The next phase, S2-P3, consists of the samples taken from units S2-SU7 (upper sample S2-193), S2-SU6, and S2-SU5, whose assemblages are similar to those of phase S2-P4 (Fig.6). Indeed, SCLEREID and SPHEROID *mixed* morphotypes are still important, with average values of 30 and 34%, respectively. The GSSCP now represent only 26% of the assemblages, still dominated by BILOBATE at 14%, and PAPILLATE are less present (> 5%) and only represented by the PAPILLATE *mixed* morphotypes. This assemblage could mark the progressive closure of the environment in connection with a change in the grass communities. This hypothesis is confirmed by the D/P index values, which are on average equal to 2. The LU index values, which are always below 0.25 (from 0.17 to 0.24), underline the presence of a grass population. During this phase, the rate of spicules gradually decreases from 26% to 16%, and diatoms are always rare (between 0% and 0.2%). These values, comparable to the S2-M3 phase, always indicate the seasonal immersion of the ecosystem.

The third phase, S2-P2, of this borehole, covers units S2-SU4 and S2-SU3 and the two basal samples of S2-SU2 (Fig.6). The SPHEROID *mixed* values are above 50% and, on average, reach 58%. SCLEREID are always present with mean values of 23%. The low cumulative GSSCP and PAPILLATE values (> 12%) allow us to affirm that for this phase, the ecosystem is covered by a *bajo* forest similar to the modern plant communities of *Chechemal*. The values taken by the D/P index in this phase are higher than 6, indicating a closed environment rich in woody plants, and the values of the LU index are higher than the threshold value of 0.25 (between 0.52 and 0.77), indicating a clearly defined environment. Spicules are, on average less represented in the assemblages with a value of 15% and a maximum at 21%, and diatoms are only present in a residual and punctual manner (between 0% and 0.3%). These data confirm the presence of a *Chechemal* forest submerged only for a short period during the year.

The terminal sample of S2-SU2 (S2-26) and the S2-SU1 unit constitute the terminal phase S2-P1 (Fig.6). Assemblages of these samples are low in SPHEROID *mixed* with an average representation of 22%, and SCLEREID values drop to 8%. The GSSCP show mean values of 24% in total and are dominated by SADDLE (11%) and RONDEL (9%) morphotypes. The PAPILLATE POLYGONAL type takes an essential part of the assemblage with values between 29 and 44%. This type of assemblage is characteristic of open *Sival* environments in Naachtun. The D/P index has values below 1 (between 0.4 and 0.7), which confirms a green environment, while the LU index has values below 0.25 (0.18), which indicates the presence of woody cover near the area. Sponge spicules show values of 30%, and diatoms are more present, with values around 10% confirming the presence of a more perennial water zone consistent with the presence of a *Sival*.

### 5.4. Statistical confirmation of phytolith assemblages

Phytolith and siliceous microfossil assemblages in the sedimentary boreholes pits and auger drilling were interpreted based on modern assemblages of these bioindicators and their respective values (Figs.3, 4, and 6). Statistical analysis is required to validate the groupings of fossil assemblages in phases. Initial analysis in a similarity cluster or HAC (Hierarchical Ascending Classification) enabled us to verify whether the ecosystem type groupings had a statistical reality. The Horn similarity index is the one that maximizes the groupings initially constructed based on assemblages (Fig.7).

**Figure 7-.**
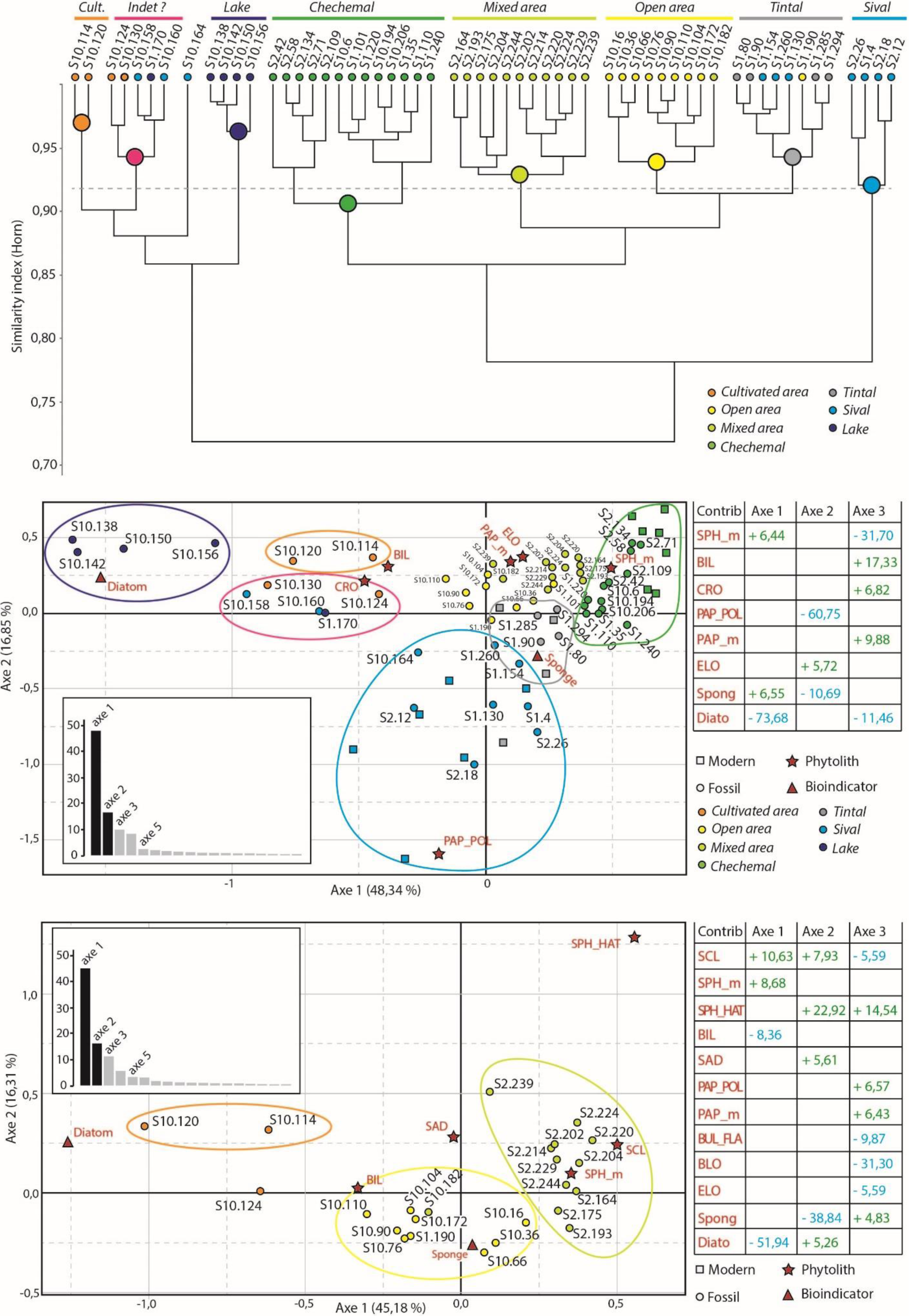
Clusters were obtained from the HAC (Horn index) of the samples of the sedimentary series. Statistical representations in the form of a CA for these same samples calibrated from modern samples taken since Testé et al. (2020).

The study of the right-hand side of the HAC (Fig.7.) allows us to validate the consistency of a certain number of sample groupings into ecosystems. All the samples initially associated with *Chechemal* phases are grouped in a single coherent cluster. Concerning the samples initially interpreted as “mixed zone” samples, all of them were finally taken from phases S2-P4 and S2-P3 of borehole pit CS1.S2c. The only exception is sample S10-182, which the HAC eventually interpreted as a typical assemblage of open plant formations. We find these ecosystems in a single cluster known as “open areas,” which finally includes only samples from borehole CS2.S10, corresponding to phases S10-P8, P7, P3, and P2. It should be noted that sample S1-190, initially interpreted as an “open area”, is not included in this cluster.

In this same right-hand part of the HAC, the surface and subsurface samples from CS2.S1 and CS1.S2c, currently located in *Sival*-type wetlands, are grouped into assemblages interpreted as *Sival*. However, a final grouping links samples initially interpreted as *Sival*, *Tintal*, and “open area” (S1-190). It can be seen that this cluster mainly concentrates samples of auger-drilling CS2.S1 from phases S1-P8, P7, P5, and P3 and that the HAC neglects the variation of these fossil assemblages. The statistical resemblance of some modern samples from *Sival* with those from *Tintal* had been highlighted in the modern reference system and explained by their ecological proximity (Testé et al., 2020). Thus, and as we will see with the forthcoming Component Analysis (CA), it would be hasty to invalidate the initial classification of these samples.

The second part of the HAC is relatively confused about sample groupings. We find a cluster comprising all the samples of phase S10-P5, which were interpreted as a deep perennial lake. On the other hand, sample S1-170 (Phase S1-P5b), also interpreted as a lake, has not been included in this cluster. The same applies to the four samples (S10-114, 120, 124, 130) initially interpreted as a cultivated area of the S10-P4 phase. Only S10-114 and S10-120 have been grouped and form an independent cluster on their own. The other two samples were eventually pooled with the S1-170 sample and the S10-P6 phase samples to form a third cluster. Only sample S10-164 is isolated in this second part of the HAC. We note that all the samples in this paragraph share the distinctive feature of having high sponge spicules and diatoms values.

Component Analysis (CA) allows the spatial location of the different groups of fossil samples identified by the HAC according to correspondence criteria. To determine whether some groupings obtained by HAC are analogous to modern environments, modern ecosystem assemblages studied at Naachtun (*Chechemal*, *Tintal*, and *Sival* - Testé et al., 2020) were added to CA but without affecting the relative contribution of the fossil samples.

Axis 1 of the turnover participates in 48% of the analysis and allows a primary organization of the samples. Diatom richness helps to attract samples with high diatom values to the negative part of the axis, while conversely, the SPHEROID *mixed* - Sponge spicule association attracts samples to the positive part of the axis. This first differentiation allows the lake samples (S10-P5) from borehole CS2-S10 to be positioned at the negative end of the axis when the *Chechemal* samples are all grouped towards the positive end of axis 1. All of *Chechemal*’s modern standards are also positioned on the axis consistently with the fossil samples.

On-axis 2, which takes part in 16% of the analysis, the samples are organized according to their richness in PAPILLATE POLYGONAL and Sponge spicules on the negative part and ELONGATE morphotypes for the positive part. Thus the samples of *Tintal* and *Sival*, both modern and fossil, are spread over the negative part of the axis. Most of the other samples are on the positive part of this axis.

The CA enables most of the groups built by the HAC to be validated. The *Chechemal*, *Tintal*, and *Sival* form three distinct point clouds calibrated by their modern analogs. The samples interpreted as lake area, cultivation area, and the indeterminate area also form three distinct point clouds. Only the samples from the mixed and open zones are too intertwined in the scatterplot to create two groups. A second CA was carried out, which included only these samples and the samples from the cultivated area.

On this second CA, which only includes fossil samples, those interpreted as cultivated areas are well differentiated on the negative part of axis 1 by the contribution of diatoms and BILOBATE. Conversely, the samples from the mixed zones of phases S2-P4 and S2-P3 are attracted by the majority contribution of SPHEROID *mixed* and SCLEREID on the positive part of the axis. The SPHEROID HAT SHAPE, SCLEREID, and SADDLE morphotypes that helps to attract these same samples to the positive portion of axis 2, thus creating a distinct grouping of the samples from phases S10-P8 P7, P3, and P2, which form another distinct group.

Carrying out a statistical CA analysis allows us to consider the HAC groupings as coherent and gives a statistical reality to our initial analyses. Thus the samples from the so-called ‘’indeterminate zone’’ are certainly to be interpreted as a wetland ecosystem different from that of modern *Sival*, tending towards lake environments. This open wetland ecosystem, dominated by the Panicoideae, is therefore found in phases S10-P6 and S10-P4 and in borehole CS2-S1 in its phase S1-P5b. The S1-190 sample, which the HAC had classified as a *Tintal* environment, finds itself, in the second CA, finally integrated into the point cloud of open areas, as it was initially classified. It is, therefore, possible to maintain this initial classification. The same is true for sample S1-130, initially classified as *Sival*, and finally finds itself classified in the middle of the CA *Sival* samples, despite being classified as *Tintal* by the HAC.

## 6. Study of past ecological variations by the malacological assemblages

### 6.1. The borehole pit CS2.S10

Malacological samples are taken from S10-SU6 to S10-SU3 and from the borehole’s surface S10-SU1 (Fig.8). The S10-SU6 corresponds to a first phase, S10-M4, which presents a significant shells concentration of around 600 to 300 individuals per liter. At these levels, the species *C. infundibulum* is dominant, with 59% to 61%. The rates of *Pyrgophorus* sp. spiny and smooth in these levels are on average less than 20% and 14%, respectively. As for the Planorbis, they do not exceed 6% of the assemblages. In these counts, the species *P. flagellata* is only represented by fragments, and the other species do not represent more than 2% of the shells in the assemblage. Finally, the Characeae show values of 4% of the assemblage. Fossil assemblages, mostly of *C. infundibulum*, have been interpreted for the Laguna coco in Belize as a phase of soil erosion linked to the drying up of water or a phase of reducing flooding processes (Bradbury et al., 1990). The sample is poor in diatoms while having sponge spicules values of around 30%, which today correspond to non-perennial wet environments. The signal provided by these assemblages of aquatic mollusks confirms the presence of a non-perennial wetland.

**Figure 8-.**
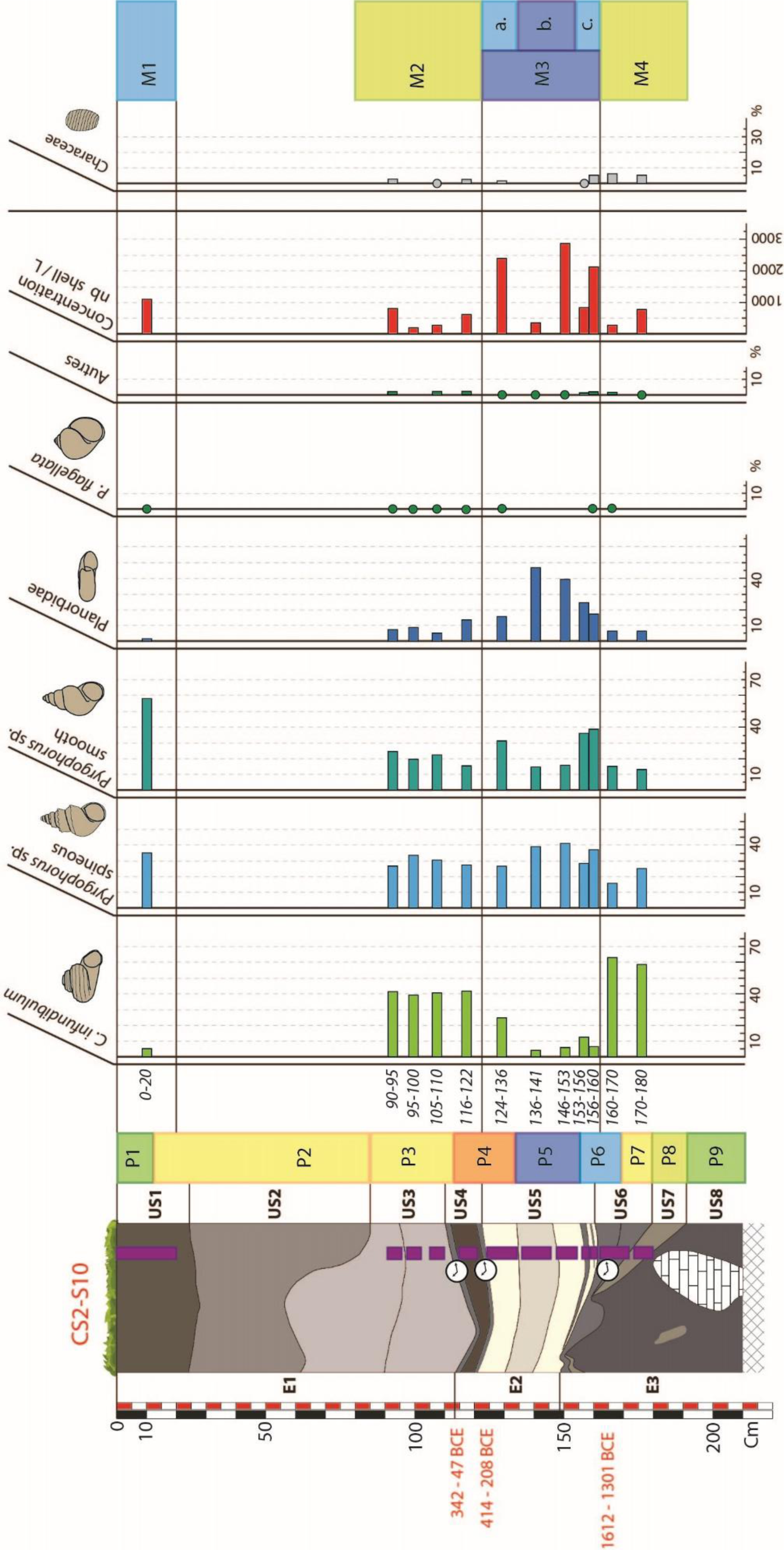
Distribution diagram of the main species of gastropods in the samples of the CS2.S10 sedimentary series, accompanied by the shell and Characeae concentrations in the samples.

The S10-SU5 is studied by five samples that form the S10-M3 phase (Fig.8). The concentration varies between 350 and 3000 shells per liter on these samples without following an evolutionary pattern. On average, the percentages of *C. infundibulum* do not exceed 6% except for a peak at 24% on the top sample (S10-124/136). The levels of *Pyrgohorus* sp. spiny remain stable between 28 and 40%. On the other hand, for smooth *Pyrgophorus* sp. shells, it is possible to follow three stages of variation in values. The first phase S10-M3c, composed of the two basal samples, shows rates of smooth *Pyrgophorus* sp. higher than 35%, then the second phase S10-M3b, composed of the two median samples, has rates decreased to 15%, finally, the third phase S10-M3a composed of the top sample (S10-124/136) shows a rate of 30%. These same three phases can also be identified for the Planorbis. The S10-M3c phase has Planorbis rates between 16% and 24%, while the S10-M3b phase doubles these values with rates between 38% and 43%. This rate decreases to 18% for the S10-M3a phase. Fragments of *P. flagellata* are still rare but are only present in the S10-M3c and S10-M3a phases. Characeae follow the same pattern and are present (between 1% and 3%) only on phases S10-M3c and S10-M3a.

Malacological work in the Laguna de cocos in Belize by Bradbury et al. (1990) was carried out on a lake much larger (600×300m) and deeper (1m50) than the *Sival* of Naachtun. The corresponding malacological assemblage, sub-surface of a borehole, shows average values of smooth *Pyrgophorus* sp. shells greater than the values of spiny *Pyrgophorus* sp., and with a rate of Planorbis shells comparable to the latter. In the deeper centimeters of the same survey, the proportions of spiny and smooth *Pyrgophorus* sp. are reversed, and the rate of Planorbs is high. This type of assemblage has been equally interpreted as a large perennial lacustrine wetland. The same assemblages have also been observed in Petén Lake with average Planorbis values comparable to the rates of *Pyrgophorus* sp. (Covich, 1976). These assemblages rich in Planorbis shells and with average levels of *Pyrgophorus* sp. are found in this S10-M3 phase and are more pronounced in the S10-M3b phase (Fig.8). The absence of *P. flagellata* in this phase S10-M3b could be interpreted as a move away from the shore (Basch, 1959), as could the absence of Characeae (Soulié-Märsche, 2002). The predominance of spiny forms of *Pyrgophorus* sp. over smooth forms could be interpreted here as the presence of a more translucent water mass (Covich, 1976) corresponding to a lake phase of the water body. Thus, phase S10-M3 can be interpreted as recording a large perennial water body that occupied the *bajo* at some point during the period ∼1450-300 BCE. Its appearance would coincide with the sudden increase in Planorbis shells (phase S10-M3c), followed by a maximum of the water body corresponding to phase S10-M3b and a decrease in the water body beginning at phase S10-M3a. This hypothesis is supported by the high concentrations of shells, the lithofacies of the carbonate deposits (sedimentology, geochemistry and micromorphology) of S10-SU5 (Castanet et al., 2022) as well as the siliceous microfossil assemblages. The phytoliths corresponding to the S10-M3c phase (S10-P6) show a more humid ecosystem, with an assembly comparable to a modern *Sival*, rest on the predominance of Poaceae and Cyperaceae and by the increase in diatoms. The samples corresponding to phase S10-M3b (S10-P5) are dominated by diatoms and suggest a larger body of water than the modern *Sival*. Finally, the siliceous microfossils corresponding to the S10-M3a phase (S10-P4) make it possible to propose a reduction of the water body with the reduction in the values of diatoms and sponge spicules.

S10-SU4 and S10-SU3 correspond to the malacological phase S10-M2, whose assemblages continue the dynamics observed in the S10-M3a phase (Fig.8). The four samples show concentration values between 200 and 700 shells per liter. During this phase, the values of *C. infundibulum* vary between 40 and 42%. *Pyrgophorus* sp. spiny levels remained relatively constant between 26 and 32%, while the smooth morphotypes fell to lower values between 17% and 24%. Planorbis are less and less represented during this phase and go from 12% to 7%. If the Characeae are still present in the assemblages, their rate does not exceed 2%. Like the S10-M4 phase, the domination of the assemblages by *C. infundibulum* shells would indicate a phase of reduction in flooding processes, which would be in continuity with a decrease in the water body observed during the S10-M3a phase. However, this signal contradicts the high levels of diatoms in the corresponding phytolith phase S10-P3, which rather indicates a perennial water body.

Finally, the surface sample of S10-SU1 corresponds to a final phase S10-M1 (Fig.8). The levels of *C. infundibulum* are 5%, while the shells of spiny and smooth *Pyrgophorus* sp. show 34% and 58%, respectively. Planorbis have residual levels of 2%, and P. flagellata are present only as a fragment. Finally, no Characeae were found at this level. This phase corresponds to a modern *bajo* forest environment near a *Sival*, i.e., an environment temporarily flooded during the rainy season and emerged the other part of the year. This dominance of smooth forms over spiny forms is interpreted as a response to conditions unfavorable to predation, such as high water turbidity (Covich, 1976; Bradbury et al., 1990). This parameter would coincide nicely with the *Sival* areas’ shallow water bodies in the Naachtun *bajo*.

### 6.2 Sedimentary series CS1.S2c

The S2-E2 basal set, studied based on core samples, did not deliver any interpretable assemblages; none of the samples had more than ten identifiable shells. This type of sampling could lead to a bias in this type of facies that a larger volume would have avoided. It is also possible that malacological concentrations are very poor at these levels. In any case, the malacological analysis of borehole CS1.S2c is based solely on the samples taken in the stratigraphic column over the S2-E1 ensemble (Fig.9).

**Figure 9-.**
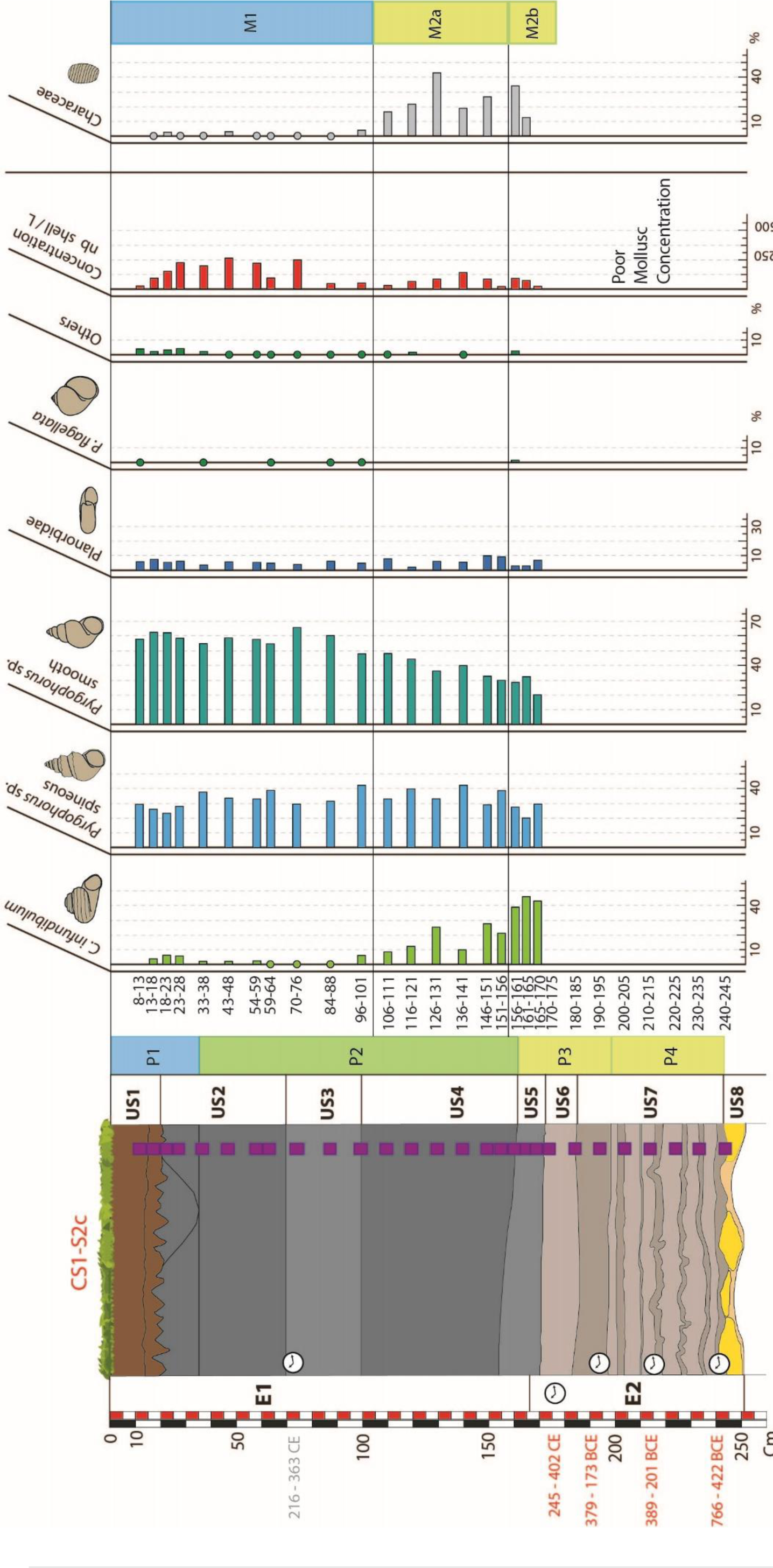
Distribution diagram of the main species of gastropods in the samples of the sedimentary series CS1.S2c, accompanied by the shell and Characeae concentrations of the samples.

Generally speaking, shell concentrations are lower in this survey than in survey CS2.S10, with average concentrations of 90 shells per Litre (Fig.9).

The S2-M2 phase is defined from the malacological assemblages corresponding to units S2-SU5 and S2-SU4 (Fig.9). The concentrations vary between 4 and 35 shells per liter without any particular pattern. On the other hand, we can see a progressive evolution of the assemblages during this phase, which makes it possible to propose two sub-phases: S2-M2b, which corresponds to S2-SU5, and S2-M2a, which corresponds to S2-SU4. The levels of *C. infundibulum* gradually increase from 44% to 9% during this S2-M2 phase, but with an average of 41% for the three samples of S2-M1and one of 17% for the six samples of S2-M2a. At the same time, we see the values of smooth *Pyrgophorus* sp. gradually increase from 20% to 49% (on average from 27% in S2-M2b to 39% in S2-M2a). Spiny *Pyrgophorus* sp. shells correspond on average to 33% of the assemblages. The values vary between 29% and 43%, except for sample S2-161/165, with a rate of 20%. As for the Planorbis represent on average 6% of the assemblages and never exceed 9%. *P. flagelatta* does not appear in the assemblages except sample S2-156/161. The other species never represent more than 2% of these assemblages. Concerning the Characeae, their concentrations vary without any particular pattern between 26% and 68%. Only two samples (S2-165/170 and S2-151/156) no present Characeae and correspond to the lowest concentration of shells in S2-SU4. In these samples, numerous bone remains of micro-vertebrates were also observed.

The dominance of *C. infundibulum* assemblages in S2-M2b has already been interpreted as an area related to fugitive water stocks (Bradbury et al., 1990). At these levels, the malacological assemblages are characterized by relatively low concentrations of shells (< 35 shells per liter) and are associated with concentrations of sponge spicules between 10 and 20% and the extreme rarity of diatoms (S2-P2). It is possible to consider this environment as the margin of a wetland drained for most of the year but marked by seasonal flooding episodes. The significant presence of Characeae in large quantities could confirm macrophyte communities associated with shallow or seasonally flooded areas. The progressive decrease in *C. infundibulum* and the increase in *Pyrgophorus* sp. in S2-M2a could suggest the gradual appearance of more sustainable wet conditions; information not found with siliceous bioindicators (Fig.6).

S2-SU3 to S2-SU1 correspond to phase S2-M1 (Fig.9). The percentages of *C. infundibulum* have decreased and represented on average 2.5% assemblages, with maximum values at 6%. The concentrations of *Pyrgophorus* sp. spiny remain stable with average values of 32% as in the previous phase. On the other hand, smooth *Pyrgophorus* sp. shells have higher average values of 58%, with peaks at 65%. Planorbis remain very poorly represented with average concentrations of 5%. The shells of *P. flagelatta* are represented mainly by fragments, and the other species are still poorly represented and less than 2% of the assemblages except for the sub-surface sample, which reaches 4%. These other species, primarily terrestrial, indeed originate from the slope. Finally, the S2-M1 phase is remarkable by the rarefaction of the Characeae, with similar values to *Sival* zones in CS2-S10.

As already shown with phase S10-M1, phase S2-M1 corresponds to installing a modern *Sival* environment (Fig.9). The wetter conditions are characterized by shell concentrations five times higher and majority assemblages of smooth and moderately prosperous *Pyrgophorus* sp. smooth and moderately rich in spiny *Pyrgophorus* sp., already found in the *Sival* areas of S10-M1. It is possible to conclude that the environment is unfavorable to predation, such as high water turbidity (Covich, 1976; Bradbury et al., 1990), a parameter consistent with the shallow, clayey waters of the *Sival* de Naachtun.

## 7. Discussion

### 7.1. Characteristics of a multiproxy study in the Maya zone

#### 7.1.1. Beneficial interactions in the multiproxy study

Phytolith assemblages enabled to interpret past plant communities based on the study of the sediments of the Naachtun *bajo*. More broadly, ecosystems and their hydrology were also approached using siliceous bioindicators, sponge spicules, and diatom frustules, based on the current reference frame produced in Naachtun (Testé et al., 2020). The freshwater mollusk assemblages have at times strengthened our interpretations (made based on the siliceous bioindicators) of the hydrological signal from boreholes CS2.S10 and CS1.S2c (Figs.8 and 9). Several concrete examples can demonstrate this.

The most explicit one concerns unit S10-US5. The mollusk assemblages rich in planorbis shells indicate a lacustrine zone (Bradbury et al., 1990). They confirmed the hypothesis of a large body of water initially based on the high proportions of diatoms. Without the mollusks, it would have been challenging to decide on the hypothesis of an increase in eutrophic conditions in the wetland leading to the formation of carbonates and a phytoplankton bloom (Rosenmeier et al., 2004).

Another correspondence exists with the *Chechemal* areas of units S10-US1 and S2-US4,3, and 2. The current sample (S10-6) from the modern *Chechemal* zone, yielded the same assemblages of siliceous phytoliths and bioindicators as the fossil samples from S2-SU3, thus interpreted as a former area of *bajo* forest. The malacological assemblage (S10-M1) of S10-SU1, which corresponds to a seasonally flooded *bajo* forest environment, is very similar to those found on S2-M2a and S2-M1. These malacological assemblages are similar to those found on the S2-SU1 unit, which corresponds to a modern *Sival* (S2-M1). Here, the comparison of all the bioindicators allows us to give an ecosystemic interpretation: the three fossil units considered at the beginning of the paragraph (S2-SU4,3, and 2) correspond to environments of *Chechemal* periodically flooded.

In general, it seems that the hydrological signal, interpreted from mollusk assemblages or concentrations of siliceous bioindicators, shows the same trends of variation. Thus on CS2.S10, these bioindicators all show a phase of increase in the water body before a phase of decrease. Conversely for CS1.S2c, the bioindicators show a progressive increase in water levels (Fig.10). Nevertheless, if these trends are similar and the correlation between gastropods and siliceous bioindicators work on certain levels, it is noted that there are some discrepancies and irresolutions that need to be discussed.

**Figure 10-.**
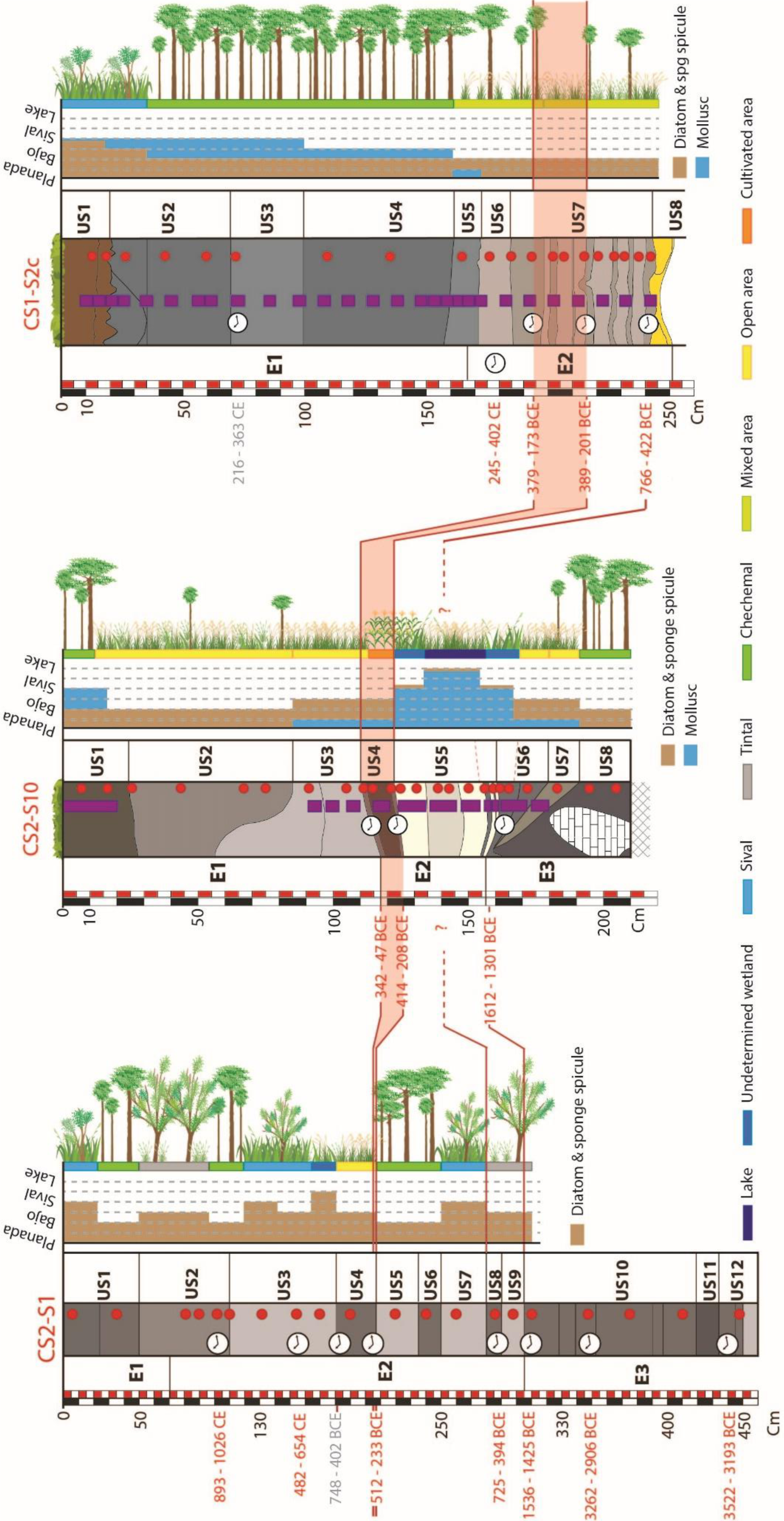
Assessment of ecosystem variations (plant communities, hydrology) calibrated from phytoliths, siliceous bioindicators, and mollusks for the three sedimentary series studied.

#### 7.1.2. Independence of ecological responses of bioindicators

The gaps in ecological responses are mainly in the CS1.S2c sedimentary series (Figs. 9, 10). In S2C-E1 assemblages, mollusks show the establishment of a *Sival*, as evidenced by the reduction of *C. infundibulum* in S2C-SU4 and the domination of *Pyrgophorus* sp. from S2C-SU3. On the other hand, Sponge spicules only indicate the presence of a *Sival* from S2C-SU2 onwards (Fig.10). This temporal shift in trends may be due to different parameters: differences in sampling, the tolerance of the organisms to ecological variations, or the taphonomy of the assemblages.

As the malacological samples were taken three years after the phytolith samples, the argilliturbation could have shifted the depths of the different sedimentary units (Beach et al., 2009). However, the short time between these two samplings and the calibration of sampling depths boreholes mitigate the effects of the process. One can above all view the sampling pitch, which is 1cm for phytoliths and siliceous bioindicators and 5cm for mollusks, and the sampling volumes are, respectively, 25cL and 5 to 10L. This very different sampling step and volume, linked to disturbed sedimentation but homogeneous facies from one SU to another, makes precision sampling difficult and may explain a shift in ecological signals over a few centimeters.

Naachtun’s modern reference (Testé et al., 2020) indicates that the dispersal of sponge spicules and fossil diatom frustules responds to an ecological differential. Sponges are organisms with fixed lifestyles, and their spicules constitute the internal ‘skeleton’ (Harrison, 1988; Frost, 2001). Thus the deposition of spicules is linked to the decomposition of the sponge and therefore to its death. Without knowledge of the porifera taxonomy of Naachtun, the distribution of spicules at scale in the *bajo* and up to seasonally flooded areas may result from two types of sponge communities. First, sponges that resist long periods of desiccation by an encystation phenomenon (Simpson & Fell, 1974) and the spicule deposits are indigenous. Second, sponges with only aquatic life cycles whose spicule deposits are transported through the *bajo* flows during flood phases. Finally, for these two cases, it is possible to imagine that the spicules deposited by aquatic communities are remobilized inside the *bajo*, as shown with the current reference system.

The modern reference system shows, in contrast to spicules, that the distribution of diatom frustules is limited to perennial water areas (Testé et al., 2020). Diatoms are mobile phytoplanktonic organisms that will follow the evolution of water masses (Smol & Stoermer, 2011). It is then customary to see a concentration of frustules in the deposits of the most perennial water zones. The microscopic size of the frustules could facilitate post-depositional transport; however, the modern datum indicates that in *bajo* environments, the transport of fossil frustules is relatively tiny: their spatial dispersion being limited to some tens of meters of permanent waters (Testé et al. 2020).

Concerning mollusks, the ecology of the Naachtun species is rather poorly described, and many data are missing. Freshwater gastropods are mobile organisms linked to restricted hydrological conditions and plant habitats (Limondin-Lozouet, 2002). However, these organisms and their clutches have numerous means of dispersal and colonization (fish, amphibian, bird, insect, drop-off) toward new environments, and they can “hibernate” when conditions are unfavorable to their survival (Dillon, 2000). It seems reasonably easy to imagine a wide dispersion of freshwater mollusk populations on the scale of a *bajo*, as shown by the homogeneous modern communities between the *Sival* and *Chechemal* areas. However, the dispersion of fossil shells on a large scale in the *bajo* by flotation seems unlikely because of the poor resistance of the shells to post-mortem transport (Limondin-Lozouet, 2002).

Finally, the life cycles and various ecological adaptations of the bioindicators used originate diachronic environmental responses. According to the taxa studied, the perception of an environmental change not express in the same spatiality or temporality. These varied responses to environmental changes are easily captured when modern taxonomic and/or ecological reference systems are available for the different bioindicators.

#### 7.1.3. Uncertainties linked to the absence of a modern reference

##### 7.1.3.1. The case of the gastropod *C. infundibulum* (Plt 5.1)

In addition to temporal shifts in the perception of ecological conditions, malacological assemblages and siliceous bioindicator concentrations can give quite different ecological interpretations of the same sample (Fig.10). Phases S10-M4, S10-M2, and S2-M2b are characteristic of this uncertainty since they indicate relatively low-water zones, close to an external aureole of *bajo* (*planada*). At the same time, spicules and diatoms are interpreted as a seasonally flooded *bajo* zone (S10-P7, S2-P3) or even a perennial water zone (S10-P6, S10-P4). There is a good chance that the poor knowledge of the ecology of *C. infundibulum* is the cause of this dissonance.

Indeed, *C. infundibulum* is nowadays only found in the Petén in a sub-fossil state (Goodrich & Van der Schalie, 1937; Covich, 1976). Today the species is found in lakes north of the Yucatán Peninsula, and its autecology has not been studied (Covich, 1976; Bradbury et al., 1990). This lack of knowledge of the species in question is felt particularly by Bradburry et al. (1990). The author interprets the high percentages of *C. infundibulum* as an indicator of freshwater, then as an indicator of a drying up of water bodies before finally considering them as indicating high hydrological levels. Here the lack of knowledge of the ecology of the species and the absence of a current reference system can easily explain the discrepancy in our paleoecological interpretations.

Therefore, it would be possible to reinterpret Fig.10 with a *Sival*-like perennial wetland, indicated by *C. infundibulum*, at least for the phase S10-P4. Nevertheless, this hydrological interpretation based on the high concentrations of *C. infundibulum* would make this time consider that the phytolith phases S10-P7 and S2-P3, which initially indicated areas with minor *bajo* flooding, are underestimated in comparison with the corresponding malacological phases. Here again, we see an irresolution due to a lack of knowledge of the hydrological conditions of life of the species *C. infundibulum*.

However, it may be possible to link variations in *C. infundibulum* to the dynamics of plant formations rather than to hydrological variations. Covich (1976) interpret the peaks of *C. infundibulum* with the leaching and erosion of the slopes of Lake Petén Itza due to deforestation, which would have increased productivity. Bradburry et al. (1990) also link high concentrations of *C. infundibulum* to palynological phases of environmental openings during the Maya occupation. Our three phases dominated by *C. infundibulum* (S10-M4, M2, and S2-M2b) correspond to very open vegetative phases of the *bajo* of Naachtun. Moreover, the Yucatan environments where they have been found are relatively less dense than those of the Petén. However, to conclude that there is a link between open vegetation, anthropic or climatic, and this gastropod species would still be premature. The uncertainties related to the ecology of *C. infundibulum* leave us another track for interpreting these data, especially for human practices signed by phytoliths.

##### 7.1.3.2. Ecological interpretation of an unknown assemblage of phytoliths

The modern open environments studied at the Naachtun site are the *Sival* areas (Testé et al., 2020). However, in these palaeoenvironmental reconstructions, many unknown assemblages are interpreted as open or semi-open vegetation (S1-P5c, S10-P7, P4, P3, P2 and S2-P4, P3 - Figs.3, 4 and 6).

In the absence of modern regional reference, botanically calibrated, that would present these assemblages, it is impossible to draw a precise interpretation of plant communities. On the other hand, index and statistical approaches can provide information on plant formations. Thus, the D:P index allowed us to validate open plant formations (S1-P5c and S10-P7, P4, P3, and P2) when the LU index confirmed that samples with D:P indices fluctuating around 1 corresponded well to mixed vegetative zones (forest with grassy undergrowth or savannah with scattered/nearby trees - S2-P4, P3). Finally, the statistical approach allowed us to confirm these hypotheses and have a reading on the most probable vegetation formations (Fig.7). Firstly it was shown that the open environment S1-P5c certainly had the same open vegetation communities as the open environment samples from borehole pit CS2.S10 (P7, P4, P3, and P2). These samples are statistically more similar to the *Sival* areas and the growing areas identified as open and humid vegetation. Conversely, the statistics confirmed that the “open” samples from borehole pit CS1.S2c (P4, P3) were more similar to the *Sival* zones than to the open zones, possibly confirming a mixed zone of herbaceous and treed vegetation.

### 7.2. Construction of the sediment – bioindicator dialogue

#### 7.2.1. Correlation between bioindicator phases and sedimentary units

When reading the phases interpreted by assemblages, one notices that they can be out of phase or in agreement with the sedimentary units. Thus, the malacological phases of CS2.S10 and CS1.S2c (Fig.8; 9), within the sampling step’s limit, corresponding to the extension of the different sedimentary units. This adequacy results from mollusks are susceptible to ecological variations in their living environment and sedimentation.

On the other hand, on the three sedimentary series CS2.S1, CS2.S10, and CS1.S2c, the phases of phytolith assemblages are often shifted from the sedimentary units. In other words, some phytolith phases straddle several sedimentary units that have different characteristics (e.g., S1-P6, S1-P4, S10-P6, S10-P4, S2-P3, S2-P2). In the case of extended phases such as S2-P3 or S2-P2, it seems that sedimentological variations do not influence the production of phytoliths and thus the vegetation interpretation. In smaller phases, straddling two very different sedimentary zones such as S10-P4 (Fig.4), it may be necessary to question the vertical migration of phytoliths (Fishkiss et al., 2010). However, in the absence of evidence to support this hypothesis - such as the enrichment of the lowest levels of phases in the smallest morphotypes - it appears here that plant communities responded to ecological variation in diachronous ways to their recording by sediments.

#### 7.2.2. Spatial perception of environmental variations

An important point to discuss is the geographical representativeness of the three sedimentary series studied with the bioindicators. They result from a selection within the campaign of 44 sedimentary auger-drillings in the closed topographic depression internal to the *bajo* and the margins of the latter. The 3 selected series are located in the following three types of current biophysical and sedimentary contexts: that of a *Sival* located in the center of a lake depression, that of a *Sival* located at the foot of a *bajo*’s slope, and that of a peripheral zone of a closed depression within a *bajo*. Although varied, these environments are not representative of all the environments of the *El Infierno bajo*. Moreover, the environments of this *bajo* are themselves very different from those of the less enclosed *bajo* south of Naachtun (Testé et al., 2020) or the *bajo* areas of the Naachtun region (Castanet et al., 2022).

Like all paleoenvironmental work, these three sedimentary series provide a fossil archive whose temporal and spatial representations are fragmentary and incomplete. These biases can be amplified according to the bioindicators used in the paleoenvironmental study. Especially in our study, phytoliths, mollusks, and others bioindicators provide an ecological signal considered as very local. While this has allowed the identification of agriculture in a specific area, it is impossible to generalize a series’s particular trend to a micro-regional scale, i.e., here to the entire *bajo*. One way to hypothesize a general trend in the Naachtun *bajo* may be to correlate all the ecological indices of the sedimentary series for the same period.

It is helpful to talk about the ecological perception of the CS2-E2 sediment body (Fig.2) to illustrate the last two biases discussed and the complementarity of sedimentary and palaeocological markers studies. Indeed, cross-interpretations of biological and sedimentary markers from the CS2.S10 and CS2.S1 series reinforce the established paleohydrological reconstructions.

For the Preclassic period, paleohydrological reconstructions of this lake depression, based on stratigraphic, sedimentological, geochemical and LiDAR data, have shown a hydrological period marked by relatively high lake levels between ∼1500 - ∼300 BCE. The lake was probably sub-perennial to perennial, from ∼1225 to ∼1050 BCE and then from ∼710 to ∼440 BCE. The latter episode (∼710-440 BCE) shows the strongest evidence of a perennial lake (Castanet et al., 2022).

The bioindicators study of CS2S10-SU5 show high diatom rates and mollusk assemblages that are characteristic of a perennial wetland that would have existed between ∼1450 and ∼300 BCE (Fig.4). On the other hand, for borehole CS2.S1, the four sedimentary units (S1-SU9, 8, 7, and 6), which form the period 1606-263 BCE, propose different bioindicators assemblages (Fig.3). S1-SU8 and S1-SU9 have an assemblage of phytoliths characteristic of a *bajo* forest like *Tintal* type. After 728-397 BCE, unit S1-SU7 shows a *Sival* assemblage, before units S1-SU6 and S1-SU5 show *Tintal*-type *bajo* forest assemblages (Fig.4). Except for the *Sival* phase (S1-SU7), most of the environments present are drained for part of the year. The paleoecological data thus reinforce the characterization of the intermittent hydrological dynamics of this lacustrine depression and, on the other hand, its perennial character, at least between ∼710 to ∼440 BCE. They also make it possible to specify that the wettest and most aquatic sedimentary environments are observed in the southern part of the lacustrine depression before ∼250 BCE while they will be observed in the median part of the lacustrine depression later.

### 7.3. Signature of agricultural practices by Phytoliths

One of the objectives of this study is to test the potential of phytoliths to characterize traces of past agriculture in *bajo* environments. The work carried out with the phytolites on fossil assemblages allows bringing answers to this question.

#### 7.3.1. CROSS Morphotype Study: Corn Cultivation in the recent Preclassic period

The richness of the CROSS morphotype in CS2.S10 and their morphometric study reveals maize as a cultivar in a *bajo* paleosol.

The dominance of ML forms over the first three phases (P1, P2, P3) corresponds to the observations made by Pearsall and Piperno (1990) and Iriarte (2003) in herbaceous ecosystems without corn. The increase and domination of L forms over the S10-P4 phase would be the direction of a cultivation phase in this area since this is observed in herbaceous communities with maize plants (Iriarte, 2003). However, it is mainly the presence of XL morphotypes that allows cultivated maize to be attested. Indeed, the presence of this type is a sufficient but not necessary condition to attest to the presence of maize in a soil sample (Pearsall and Piperno, 1990; Iriarte, 2003). Samples from S10-SU3 and S10-SU2 show numerous ML morphotypes and no evidence of maize indicator XL morphotypes. This reduction in size and percentage of CROSS would be consistent with the non-use of this area for food production after the S10-P4 agricultural phase (Fig.6).

The pedo-sedimentary unit S10-SU4 has the characteristics of a cultivated paleosol. It shows traces of pedogenesis and plants using the C4 pathway (bioturbation, relatively increased organic carbon content, ^13^C enrichment of organic matter; Castanet et al., 2022) and contains phytolith bioindicators of a crop such as corn. This paleosol shows traces of redox, which validates the paleoecological interpretation made above. Indeed, this SU includes, on the one hand, malacological assemblages indicating a seasonally drained zone and, on the other hand, spicules, diatoms, and Characeae indicating flood phases or the proximity of an aquatic environment.

These paleoecological reconstructions demonstrate the establishment of agricultural practices in the *El Infierno bajo*. They provide partial information on the cultivation systems established in an extensive or intensive context, based at least on maize, implanted near wetlands, and this, since the recent Preclassic. The first archaeological building structures on the site are also dated to this period (Hiquet, 2020).

These results may inform the intensive agricultural practices carried out within the *El Infierno bajo* and other *bajo*s in the Naachtun sector as recently proposed based on the LIDAR studies conducted in the region. Numerous hydraulic and agrarian structures known as wetland features have been described and studied (Canuto et al., 2018; Castanet et al., 2019; 2022).

#### 7.3.2. SPHEROID FACETATE: traces of agriculture for the archaic period?

The S10-P9 phase shows SPHEROID FACETATE values of 4% (Fig.5). These phytoliths are produced in small quantities by Cucurbitaceae (Piperno et al., 2000). These levels are dated before 1600 years BCE. Gourds are already cultivated in Mesoamerica at this period (Colunga-García-Marín and Zizumbo-Villarreal, 2004), so it would be possible to consider these traces as witnesses of very early agriculture established in the Naachtun sector during the Archaic period.

However, observation of the assemblages shows that for each forest zone of the western section boreholes, the SPHEROID *mixed* are always associated with the residual SPHEROID FACETATE. In fact, in the areas of the *bajo* forest near the *Sival* zones, there are numerous plants of *Cucurbita radicans*, a vine belonging to the Cucurbitaceae and which produces small gourds. Cucurbitaceae morphotypes have also been found in modern *bajo* environments in Naachtun (Testé et al., 2020). In our case, it is more likely that these phytoliths originated from wild and local populations of Curcubitaceae than from an early agricultural phenomenon in Naachtun. This interpretation also suggests that the discovery of squash phytoliths is not a systematic condition for its cultivation.

### 7.4. Contribution of bioindicators to an environmental history of Naachtun

Most of the established paleoecological reconstructions concern the periods from the ancient Preclassic to the early Classic periods. The current archaeological knowledge of Naachtun is based on work that has provided information on the period between the end of the late Preclassic and the Postclassic (Nondeodeo, 2013; Sion, 2016; Hiquet 2020). For lack of being able to reconstruct fully and with good resolution the interactions between the population of the Maya city of the Classic period and its environment during the apogee of Naachtun, we obtain results and discuss socio-environmental interactions relating to the occupation of the area during the Preclassic, before the emergence of the city.

Despite the early presence of Cucurbitaceae phytoliths in the basal assemblages of CS2.S10, the hypothesis of archaic or ancient Preclassic agriculture at Naachtun has not been retained in this work for the time being. On the other hand, in this same series, we note the early opening, dated at least to the ancient Preclassic, potentially older, of this area of the *bajo* with the appearance of Poaceae communities. The lack of palaeoecological data (CS2.S1) or sedimentary archives (CS1.S2c) for this period prevents discussion of a generalized opening of the *bajo* zone. Nevertheless, the transformation of hydrosedimentary dynamics identified in this lake depression around 1500 BCE have been interpreted as a response of the lake hydrosystem and morphosedimentary system to a watershed-wide anthropogenic impacts as opening of the vegetation cover (HP3 and ESTP3a, Castanet et al., 2022). Vegetation opening during the archaic or early preclassic period has been revealed by bioindicators of many wetlands in the CMLs (Islebe et al., 1996; Curtis et al., 1998; Wahl et al., 2006; 2007; 2013; Carozza et al., 2007; Luzzader-Beach et al., 2016). Although the drier climate of the early prelassic (Douglas et al., 2016; Rosenmeier et al., 2016) has been questioned concerning the extension of herbaceous zones, their diachronous development, and the presence of cultivars (Wahl et al., 2006; Wahl et al., 2013) instead leads the authors to support the thesis of anthropogenic impacts in different areas of the CMLs region.

On all Preclassic levels of CS2.S10, data on hydrological bioindicators indicate the presence of a wetland (Fig.5;10). This perennial wetland is interpreted as a lake with intermittent dynamics, which have been perennial at least during a part of the middle Preclassic if the dating of the wettest phase of CS2.S1, based on bioindicators, is believed. These results are broadly in line with the paleohydrological and geoarchaeological work based on the study of stratigraphy, sedimentary and geochemistery markers carried out on these same archives (Castanet et al., 2016; 2022). This hydrological period of perennial lake dynamics corresponds to a period of increased precipitation or decreased evaporation (mid-recent Preclassic) observed in several places in the CMLs. We note studies about the δ^18^O of Lakes Salpetén and Puerto Arturo (Douglas et al., 2015; 2016; Rosenmeier et al., 2016), the diatoms of Lake Tuspan (Nooren et al., 2018), the δ^18^O of Macal chasm speleothems (Webster et al., 2007) or based on spatialized palynological reconstructions of the region (Carrillo-Bastos et al., 2012). The presence of a large water-body, as is the case for many Maya sites (Tikal, El Mirador, La Joyanca, Yaxha), could have favored an early preclassic Maya settlement on its territory.

The earliest archaeological evidence of occupation of the Naachtun microregion dates to the Middle Preclassic period around the 6^th^ century BCE (Nondédéo et al., in press). In the centre of Naachtun, the earliest archaeological evidence of occupation is dated to the Late Preclassic. This archaeological evidence is consistent with the identification of cultivated soil in CS2.S10, attested by cultivator phytoliths, δ^13^C and micromorphology, thus demonstrating the cultivation of maize in the *bajo* wetlands during this period (412-49 BCE). It also adds an element in favor of *bajos* in the food production of Maya societies (Turner and Harrison, 1981; Pohl, 1990; Dunning et al., 1998). The correlated dates around the period 400 - 200 years BCE of the three sedimentary series over this period (CS1.S2C, CS2.S1, and CS2.S10) show rich assemblages of herbaceous Panicoideae (Fig.10). These assemblages presuppose a substantial opening of the environment in the *bajo* or at least a herbaceous component of the ecosystems strong enough to suggest a general opening of the *bajo*. This phenomenon could result from intensive agricultural practices within the *bajo*, as suggested by increasing the phytoliths associated with the Panicoideae. An episode marked by increased erosion and transfer of sediments from the uplands to the lowlands has been identified from the study of the sedimentary series of the *bajo El Infierno* from ∼1500 BCE. This major episode of Maya clays deposits has been interpreted as a response of the morphosedimentary system to deforestation and agrarian practices developed on the uplands (ESTP3a, Castanet et al., 2022). The practice of agriculture within *bajos* of CMLs during this period has also been documented by work on CMLs wetland bioindicators, including maize pollen (Pohl et al., 1996 ; Wahl, 2006; 2013; Carozza et al., 2007; Krause, 2019) but also by the geomorphology and geoarchaeology that revealed cultivated soils (Beach et al., 2009; Dunning et al., 2019) or agrarian structures such as mounds or raised fields (Dunning et al., 2019; Beach et al., 2019). Furthermore, these same types of structures have also been updated by LIDAR techniques in the *bajos* of the CMLs, especially in Naachtun (Canuto et al., 2018; Castanet et al., 2019). Bradbury et al. (1990) propose that an increase in hydrological levels in Belize (Laguna Cocos) during the Preclassic period led the Maya to use the *bajo* and wetlands for agriculture. In the Elevated Interior Region, we find this double hydrological and agrarian signal on the paleosol of CS2.S10 with high concentrations of diatoms and sponge spicules. This hypothesis also coincides with the regional paleoclimatological framework, which indicates a wet period in the middle and recent Preclassic period (Webster et al., 2007; Carrillo-bastos et al., 2012; Douglas et al., 2015; 2016; Rosenmeier et al., 2016; Nooren et al., 2018).

The sedimentary environments of these Preclassic periods are diversified. On CS2.S10 and CS1.S2c, these sedimentary environments correspond to the S10-E2 and S2-E2 sediment bodies (Fig.10). The S10-E1 and S2-E1 sediment bodies are thick, mainly weakly carbonated silty clay and argilliturbated. The age models established for these series allowed to assign an age of the Classic period to the lower part of these deposits (Castanet et al., 2022). SU5 to SU3 of the CS2.S1 series corresponds to levels of the Classic. The study of these levels shows that the plant communities appear to be relatively different from one series to another, indicating a mosaic of landscape in the *bajo* and a diversity of wetland management during the Classic period.

Borehole pit CS2.S10 shows open environments rich in Panicoideae (Fig.4) along E1 sediment body. Maintaining an open formation in a *bajo* zone that was only flooded seasonally and is now covered by forest when it is abandoned, suggests that this zone was still under human influence, which remain to be specified. In CS1.S2c, phytoliths and mollusks indicate a closure of the ecosystems at the beginning of the Classic period and the presence of a *bajo* forest plant community (Fig.6). The presence of this type of forest in the *bajo* is consistent with the charcoal analysis results for the Naachtun area, which show the use of many typical secondary or *bajo* woods during the ancient and recent Classic period (Dussol, 2017). There is likely a change in land use in this part of the *bajo* that could possibly result from agricultural abandonment in the CS1.S2c area. This abandonment, and its possible generalization to the entire *bajo*, would raise questions about the reorientation of food production in the growing demographic context of the Classic period (Hiquet, 2020).

Finally, on CS2.S1, the Classic levels are marked by an alternation of *bajo* forest and *Sival* zones at least since 433-651 CE. The Naachtun *Sival* wetland as we know it today (centered on CS2.S1 - Fig.1) therefore seems to have been established during the Classic period, at the same time as the CS2.S10 series seems to be less and less subject to the action of a body of water (Fig.10). This is largely the result of the transformation of the intermittent lake depression initiated in response to detrital inputs that occurred between 1500 and 100 BCE and continued between 250 and 1000 CE (ESTP3 and ESTP5, Castanet et al, 2022).

This mosaic of landscapes (savannah, forest, wetlands) allows us to doubt climate change action during the Classic period to explain such substantial variations on such a small scale. On the other hand, demographic and climatological data show us that the transition between the Preclassic and Classic (100 - 300 CE) is marked by several events. In Naachtun, the Classic/Preclassic transition is part of a demographic process in which non-natural population growth is observed and could result from the abandonment of the large surrounding cities. These populations could very well have arrived with other or unused practices in Naachtun. Finally, a drying up of the climate is noted during this same transition period, End of the Late Preclassic / beginning of the Early Classic (Douglass et al., 2015, 2016; Rosenmeier et al., 2016). This drought could also have pushed the local Maya populations, who used the *bajos* for their agricultural production, to diversify their subsistence practices. However, these hypotheses, based only on the naturalistic inheritance of bioindicator assemblages and geomorphology will need to be confronted with new geoarchaeological data of this site, particularly from LiDAR (Canuto et al., 2018; Castanet et al., 2019) and from the study of intrasite paleosols (Castanet al., 2016).

## 8. Conclusion

Palaeoecological reconstructions based on bioindicators are difficult to carry out in Maya areas and have often been restricted to perennial water zones. This study shows that phytoliths can be used in the sediments of a seasonally drained wetland to reconstruct ancient ecosystems. The existence of a repository of modern ecosystem phytoliths was a determining factor in interpreting fossil data, as was their statistical treatment, index approaches, or morphometric study of particular phytoliths. The complementary use of bioindicators such as the fossil shells of gastropods and sponge spicules and diatoms concentrations made it possible to confirm specific hypotheses on the hydrological state of the wetland that was not resolved with phytoliths alone. What emerges from this multiproxy approach is that the cross-referencing of ecological signals makes it possible to balance out all the taxonomic biases and gaps in ecological knowledge specific to each bioindicator. It has been possible to interpret the variations in the assemblages of phytoliths, siliceous bioindicators, and fossil mollusks in terms of ecosystem change over time. Moreover, paleoecological reconstructions based on bioindicators reinforce those based on sedimentary and geochemical proxies.

The earliest records of phytoliths show a *bajo* forest that gradually opened up at least before 1600 BCE, most certainly during the Archaic period. For the hydrological period marked by relatively high lake levels between ∼1500 and ∼300 BCE (Castanet et al., 2022), the study of siliceous bioindicators and mollusks show that perennial hydrological dynamics was particularly pronounced from ∼700 BCE. In the Late Preclassic period, more specifically between 400 and 200 BCE, there are the first traces of agriculture in the wetland in the form of maize phytoliths and paleopedological indications. During this same period, it is above all the presence of herbaceous formations that is noted, as attested by phytoliths or mollusks on the three sedimentary series. This opening of the environment coincides with the first archaeological traces of occupation identified to date for the edge of the *bajo El Infierno*.

The end of the Late Preclassic and beginning of the Early Classic was marked by a demographic change associated with the growth of the Naachtun centre and a drying up of the climate. For the classic period, despite the sediment archives and the calibrated age models, the local spatial expression of the used bioindicators did not reveal major ecological patterns in the bajo, but rather a landscape mosaic alternating between herbaceous formations, palustrine environments, and forests. If this local landscape diversity can result from numerous adaptations of the inhabitants of Naachtun to the social and ecological contexts, it demonstrates above all the need to carry out multi-spatial approaches in paleo-environmental reconstructions in Maya areas to understand the local occupation of a territory. These reconstructions based on biological indicators thus strengthen the knowledge of the dynamics of the Maya socioecosystem that persisted in this microregion for at least 2,500 years, during the Preclassic and Classic periods (Castanet et al. 2021).

Although the objectives of this study have, for the most part, been validated - i.e., the reconstitution of palaeoecosystems using assemblages of phytoliths and related bioindicators - there are still biases that need to be mitigated to make this approach more utter. The first bias that emerges is the presence of phytolith assemblages, unrecorded in our modern reference (Testé et al., 2020), which thus reflect ecosystems not currently known in Naachtun. The key to interpreting these new assemblages is to complete the modern reference systems, especially by studying Maya agrosystems as recently undertaken by our research team (PAYAMA project; Dussol et al., 2021). Finally, the second central bias concerns the use of mollusks in wetland sediments. Although the ecologies of the region’s terrestrial mollusks are well known (Dourson & Caldwell, 2018), this is not the case for freshwater mollusks. That is particularly true for the ecology of the *Cochliopina infundibulum* species, one of the species with the most marked variations in the sediments of Naachtun. This lack of knowledge of the ecological valences of gastropod species and the lack of information on modern biocenoses limits the interpretation of fossil data. Ecological work, in the form of an ecological reference system of assemblages, on the malacological communities of the Petén ecosystems, would initially provide better information on the living conditions of the freshwater gastropods found and improve the paleoecological interpretation of these data.

## 9. Acknowledgement

This research is supported by the Naachtun Project 2010-2018 (Dir. Ph. Nondédéo), the HydroAgro Project (coord. E. Lemonnier and C. Castanet), the PAYAMA Project (coord. A. Garnier and E. Garnier), the LabEx DynamiTe (ANR-11-LABX-0046), the Ministry of Foreign Affairs (France), the Pacunam Foundation, the Perenco company (Guatemala), the CNRS (France) and the Université de Paris 1 Panthéon Sorbonne. This research is supported by the institutional support of the Centre for Mexican and Central American Studies (Guatemala).

This publication has received financial support from the Treilles Foundation. The Foundation des treilles was created by Anne Gruner Schlumberger and aimed, in particular, to open and nurture dialogue between science and the arts in order to advance contemporary creation and research. It also welcomes researchers and writers in the field of Treilles (Var) - www.les-treilles.com.

Finally, we pay tribute to our precious botanist and friend, *Enecon Abelino Oxlaj*, who unfortunately left us last year. His fieldwork was crucial in the realization of our research. Our thoughts are with his family and friends.

## 10. Compliance with Ethical Standards

### Funding

Marc TESTÉ has received research grants from LabEx DynamiTe and from University Paris 1 Panthéon Sorbonne. For the finalization of this article, he also received the support of the Fondation des Treilles. Cyril CASTANET has received research grants from University Paris 1 Panthéon Sorbonne. Aline GARNIER has received research grants from LabEx DynamiTe. Nicole LIMONDIN-LOZOUET declares that she has no conflict of interest. Louise PURDUE declares that she has no conflict of interest. Eva LEMONNIER has received research grants from University Paris 1 Panthéon Sorbonne and from LabEx Dynamite. Sébastien KERDREUX declares that he has no conflict of interest. Philippe NONDÉDÉO leads the Naachtun project and he receives research funds from Ministry of Foreign Affairs (France), from the Pacunam Foundation and from the Perenco company (Guatemala).

### Ethical approval

This article does not contain any studies with human participants or animals performed by any of the authors.

